# Analysis of the peptide helicity using an ensemble spectroscopic model with re-calibrated parameters

**DOI:** 10.1101/2023.10.25.563921

**Authors:** Uroš Zavrtanik, Jurij Lah, San Hadži

## Abstract

The α-helix is the fundamental building block of protein structure. Understanding the physical principles that determine how a specific amino acid sequence defines helicity is a key step towards elucidating the sequence-structure relationship. An established method for quantitation of helix content using circular dichroism (CD) relies on the linear spectroscopic model. In this model, the helix length-correction is not applied to each ensemble conformer individually, rather an average value is assumed for all conformers. Here we assess the validity of this approximation and introduce a more physically realistic ensemble-based analysis of the CD signal. We find that the linear model tends to underestimate peptide helicity, with the difference depending on the ensemble composition. Using a CD dataset covering a broad range of helicities, we re-calibrate spectroscopic parameters (helix and coil baselines) and determine helix-coil parameters for a set of alanine-rich peptides. Our results show that the ensemble model can leverage the small spectroscopic differences between peptide conformers, enabling it to extract more information from the experimental data. We show that some previously poorly-defined quantities, such as helix nucleation constant and heat capacity change associated with helix folding, can be reliably determined using ensemble model. Overall, the presented ensemble-based treatment of the CD signal, together with the re-calibrated values of the spectroscopic baseline parameters, provides a coherent framework for the analysis of the peptide helix content.

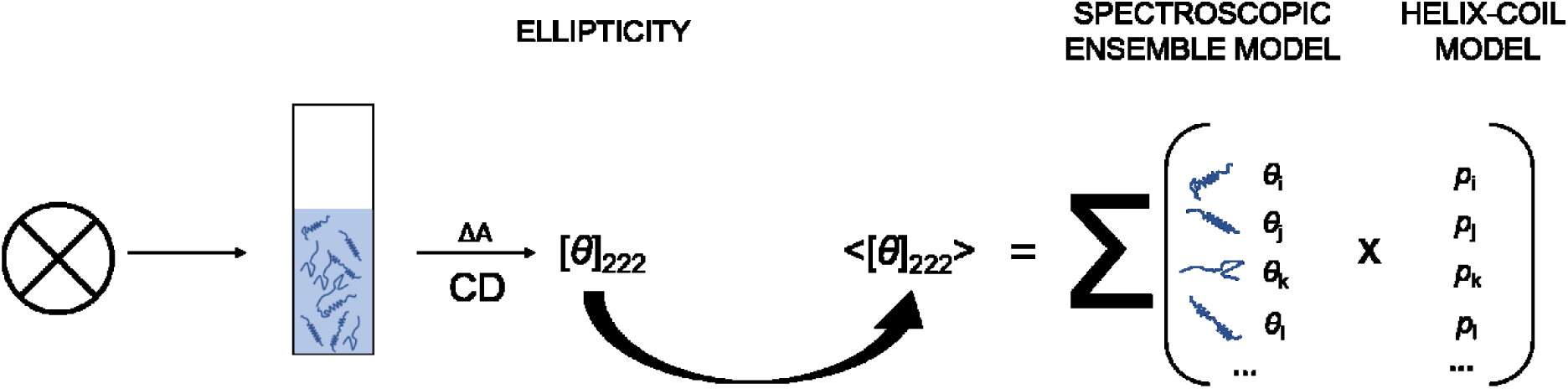

## Introduction

Circular dichroism (CD) is a powerful spectroscopic technique for quantification of the secondary structure in proteins and peptides. The CD originates from differential absorption of left- and right-handed circularly polarized light, and the difference is typically measured as the ellipticity of the transmitted light. The primary chromophore in proteins is the peptide bond which exhibits strong absorption in the far-UV region. The shape of the CD spectrum is determined by two key transitions: the n-π* and π-π* transitions, both of which are sensitive to the backbone torsion angels (secondary structure) [1]–[3]. For helical peptides, the CD spectrum has two minima at 222 and 208 nm and a maximum at 190 nm, while peptides in coil conformation have a characteristic negative peak at 200 nm. The minimum at 222 nm is due to the n-π* transition, which is particularly pronounced in the α-helix and can be used to quantify the helix content. The transition from helix to coil exhibits an isodichroic point at 203 nm, indicating that a single chromophore (peptide unit) can populate one of the two states, helix or coil [1], [3]. For a given peptide, the measured mean molar ellipticity at 222 nm, [*θ*]_222_, is the sum of the spectroscopic contributions of all peptide units in helix and coil states. Furthermore, in solution a peptide can sample an ensemble of *M* different conformers, each with a unique sequence distribution of helical and coil units. The total [*θ*]_222_ signal for an ensemble of *M* conformers can be given as follows:

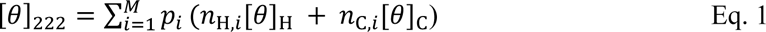

Here, each *i*-th conformer has *n*_H,*i*_ and *n*_C,i_ peptide units in the helix and coil states. These contribute [*θ*]_H_ and [*θ*]_C_ to the total CD signal, which are the molar ellipticities of helix and coil. The signal contribution of each *i*-th conformer is weighted by its ensemble probability *p*_i_, thus the total signal measured [*θ*]_222_ is a sum of the probability-weighted contributions of all *M* conformers in the ensemble. To determine the ensemble composition (probability of each conformer) from the measured [*θ*]_222_, one needs to know the values of the helix and coil molar ellipticity ([*θ*]_H_ and [*θ*]_C_), commonly referred to as helix and coil baselines.

The coil baseline, [*θ*]_C_, has been estimated experimentally using model peptides in which all units were assumed to adopt the coil conformation. Short peptides or high-temperature data presumably satisfy this requirement, and the obtained [*θ*]_C_ ranges from 600 to 2200 deg cm^2^ dmol^-1^ at 0°C [4], [5] (all values in the manuscript are per mole of peptide unit). The coil ellipticity changes with temperature and is considered additive for a given sequence (independent of peptide length). The estimation of the helix baseline [*θ*]_H_ is more difficult, and its values are still debated [6]. The reason for this is twofold. First, it is difficult to obtain systems in which all peptide units would adopt only the helical conformation due to low cooperativity of the helix-coil transition. Second, the magnitude of peptide helix ellipticity strongly depends on the length of the helical segment to which it belongs [7], [8]. This length-dependent effect is described by an empirical end-effect parameter *k*:

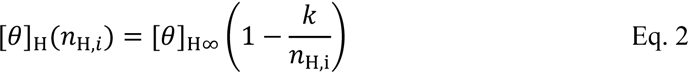

where, [*θ*]_H_ (*n*_H,i_) corresponds to the molar ellipticity of a peptide unit in the helix with total of *n*_H,i_ helical units. The [*θ*]_H**∞**_ is the molar ellipticity of an indefinite helix, and the right-hand term is the correction for the end-effect scaled by parameter *k*. For a sufficiently long helix (*n*_H,*i*_ >> *k*), the correction becomes insignificant and [*θ*]_H_ ≈ [*θ*]_H_**_∞_**. However, peptide units in shorter helices have a lower (less intense) ellipticity, which depends nonlinearly on *n*_H_ (Eq. 2). Therefore, to calculate the [*θ*]_H_ contribution for a peptide, the length of the helix (*n*_H_) to which the peptide belongs and the spectroscopic parameters [*θ*]_H_**_∞_** and *k* must be known. Several theoretical and experimental approaches have been used to estimate the helix baseline parameters, but there is no consensus on these values. The proposed values range from −35×10^3^ to −44×10^3^ deg cm^2^ dmol^-1^ for [*θ*]_H_**_∞_** and from 2 to 4 for *k*, but values up to *k* = 6 have also been proposed [9].

What is the molecular origin behind the length-dependence of the helix CD signal? In 1941, Kauzmann and Eyring argued that helical peptide units experience different restriction from the intrapeptide hydrogen bonds [10]. The three peptide units at the ends of the helix have single hydrogen bond, whereas the units inside the helix are double bonded to *i*-3 and *i*+3 peptide groups (**Figure 1**). Double-bonding imposes greater constraint on the O=C-N plane, which enhances the signal compared to the six residues at the helix ends. This idea was supported at the time by the difference in optical activity of open chains and closed rings of polyhydroxy alcohols [10]. This line of thought was taken up by Shalongo and Stellewagen, who developed a dichroic spectroscopic model to explain the helix signal length-dependence [11]. In this model, two spectroscopic contributions are explicitly assigned to the single- and double-bonded peptide units (**Figure 1**). Double-bonded units in the helix center correspond to the situation in an infinite helix and are therefore assigned [*θ*]_H∞_, while the single-bonded units at helix ends have ellipticity [*θ*]_H1_. The empirical end-effect parameter *k* can be expressed as a function of [*θ*]_H1_ and [*θ*]_H∞_ [11]:

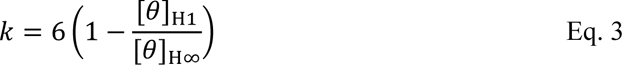

**Figure 1.**
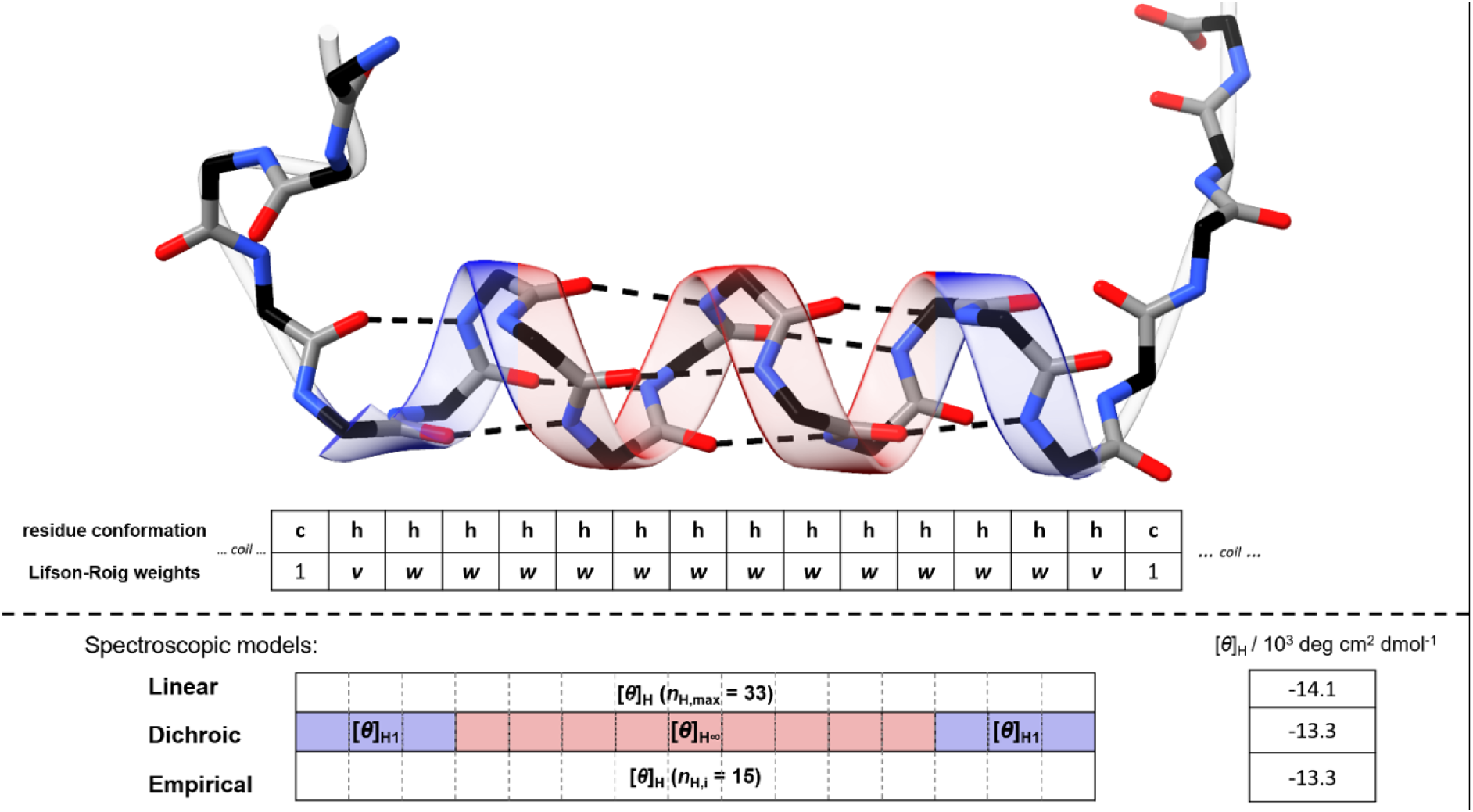
Assignation of signal contributions by different spectroscopic models. Upper panel shows a helix segment spanning *n*_H_ =15 helical units as part of a peptide with a total of *N*_pep_ = 33 units (only some of the units neighboring the helix are shown for clarity). Coil units are shaded in gray, helical ones are in red and blue, depending if they are double or single hydrogen bonded. Upper table shows residues definitions (coil or helix conformation) and the corresponding LR statistical weights. Lower table shows peptide units and the corresponding ellipticity contributions in linear, dichroic and empirical models. Values listed on the right illustrate the differences in ellipticity calculated with three spectroscopic models using parameters [*θ*]_H∞_ = −40×10^3^ deg cm^2^ dmol^-1^, *k* = 4, *N*_pep_ = 33 and [*θ*]_H1_ calculated by Eq. 3.

In the dichroic model, peptide chromophores are counted individually. The nonlinear [*θ*]_H_ (*n*_H_) dependence is due to the changes in the ratio of single and double-bonded units. In sum, there are two approaches to describing the length-dependence of the helix CD signal. One approach, the empirical model, treats the length-dependence at the level of the entire helix and yields a continuous signal dependence defined by the parameter *k* (Eq. 2). The second approach, the dichroic model, operates at the level of individual peptide units (chromophore). The signal contributions are treated as discrete units and the end-effect is explicitly assigned to the terminal peptide units. As we show below, they give very similar results and differ only when the helical segments become very short. Both models consider the spectroscopic signal as an ensemble property; the value of the helix signal depends on each conformer state; therefore, we refer to these two models as ensemble models.

Interestingly, the distinction between the two ensemble models is obscured by the third, most widely-used method for calculating the helix content. The so-called linear model uses the empirical length-correction (Eq. 2), however the same correction is applied to all conformers in the ensemble, regardless of the actual lengths of their helical segments. The helix ellipticity is corrected by assuming a maximal helix length for all conformers, i.e. instead of [*θ*]_H_(*n*_H,*i*_), [*θ*]_H_(*n*_H,max_) is used in Eq. 1 (**Figure 1**). This simplifies Eq. 1, since the sum over the probability-weighted number of helix contributions is equal to the product of [*θ*]_H_ (*n*_H,max_) and the average number of helix residues <*n*_H_>, which can be easily derived from the partition function [12]. As such the linear approximation essentially enables to calculate helix content from [*θ*]_222_ without knowing the ensemble composition, which is not possible when [*θ*]_H_ is evaluated separately for each conformer, as in ensemble models. For this reason, virtually all studies use the linear model to estimate the helix content from CD data. Ensemble spectroscopic models are rarely used, in part because the usual matrix approaches for calculation of partition function are not well-suited for applying individual, per-conformer end-effect corrections. This is because the conventional helix-coil matrix format yields the expanded matrix product (partition function polynomial), where spectroscopically distinct states are grouped in the same terms. To circumvent the problem of mixed matrix terms, Gans et al. used an iterative enumeration algorithm to calculate the peptide ensemble [9]. This approach has the advantage that individual conformers can be better tracked and spectroscopic contributions were then assigned using the empirical model (Eq. 2) [9]. Later, Shalongo and Stellwagen presented a critique of the linear approximation model [11] and used the enumeration algorithm to assign contributions using the dichroic model. Significant differences were found compared to the linear model, but the study used different definitions of the chromophore unit between the models, making direct comparison difficult. While the linear approximation is reasonable for highly helical ensembles (<*n*_H_> approaching *n*_H,max_), it is not entirely clear to what extant the approximation also applies to less helical ensembles.

The analysis of the CD signal in helical peptides concerns two main issues: i) the accuracy of the spectroscopic parameters and ii) the use of an appropriate model to assign the spectroscopic contributions. Despite that CD is the most common method for quantifying helical content, neither of these issues has been satisfactorily resolved. In the linear model, the paradoxical situation arises since the system is modeled as an ensemble at the molecular level, but from a spectroscopic point the ensemble character is partially lost. This raises several questions. What precisely are the differences between the three spectroscopic models (empirical, dichroic, linear)? Namely, how much would the [*θ*]_222_ values calculated with different models differ for a given ensemble? The inverse question is also relevant: Do we get different estimates of the helix-coil parameters from the [*θ*]_222_ data depending on the choice of spectroscopic model? In the past, much work has been dedicated to improving helix-coil models and determine their parameters, but less attention has been paid to the spectroscopic models that link the measured signal to the molecular ensemble. Moreover, the spectroscopic parameters differ between studies. Interestingly, some typical values used as coil and helix baselines appear to be based on unpublished data or personal communication [4]. Given the spread in these numbers, there seems to be no consensus on the spectroscopic baseline parameter values.

Here, we first develop a new matrix-based algorithm to compute the peptide partition function. The expanded matrix product gives individual terms in the partition function polynomial for different helical conformers allowing simple application of helix length-correction to each conformer. We then evaluate the differences between the spectroscopic models and observe that the linear model tends to overestimate the CD signal and underestimate the peptide helicity. The difference depends on the ensemble composition and increases with the number of intermediate-sized helix conformers and double-helix conformers. To determine the spectroscopic baseline parameters in consistent manner, we assembled a diverse dataset of measured ellipticities ranging from all-helix peptide systems to the alanine peptide series (AAKAA)_n_-GY and a short peptide with low helix content. We then applied a global fit followed by Bayesian inference of the model parameters using Markov Chain Monte Carlo to simultaneously determine the helix-coil and spectroscopic baseline parameters. The empirical model yields a noticeably poorer quality of fit, while dichroic and linear models perform almost identically. However, compared to the linear model, the dichroic model exhibits significantly better parameter stability and lower parameter cross-correlations and uncertainties. Therefore, the dichroic model is not only more physically realistic, but also able to extract more information from the data because it leverages subtle spectroscopic differences in the peptide conformers. We suggest new values for spectroscopic baselines to be used for peptide helicity estimation and provide a computer program for calculating helicity from the CD data using dichroic ensemble model.

## Results

### Linear, empirical and dichroic models assign different ellipticity contributions

To understand the differences between the spectroscopic models, we calculated the molar helix ellipticity [*θ*]_H_ for the individual conformers which differ in number of helical peptide units *n*_H_. As *n*_H_ increases, the ellipticity [*θ*]_H_ increases, i.e., it becomes more negative (**Figure 2A**). This dependence is linear for the linear model but nonlinear for both ensemble models. For the maximal-length helix segment (*n*_H_ = 33), all three models converge to the identical value [*θ*]_H_ (*n*_H,max_). In the linear model, this value is used as a constant per-peptide value for all other, less-then-maximal helical segments, explaining why signal changes linearly with the number of helical units (**Figure 2A**). In contrast, in both ensemble models, the helix ellipticity is re-evaluated for each *n*_H_ (Eqs. 2, 3), leading to the nonlinear dependence and overall lower (less negative) [*θ*]_H_(*n*_H,i_) contributions (**Figure 2A**). The linear model therefore tends to systematically overestimate the CD signal or, for a given signal, underestimate the true helicity. At the level of individual conformers, this difference is most pronounced for the shorter helices (by about 1300 deg cm^2^ dmol^-1^ per peptide (**Figure 2A, inset**). Overall, the linear model gives an “averaged” picture of helix ellipticity and is insensitive to differences between short, intermediate and long helical segments.

**Figure 2.**
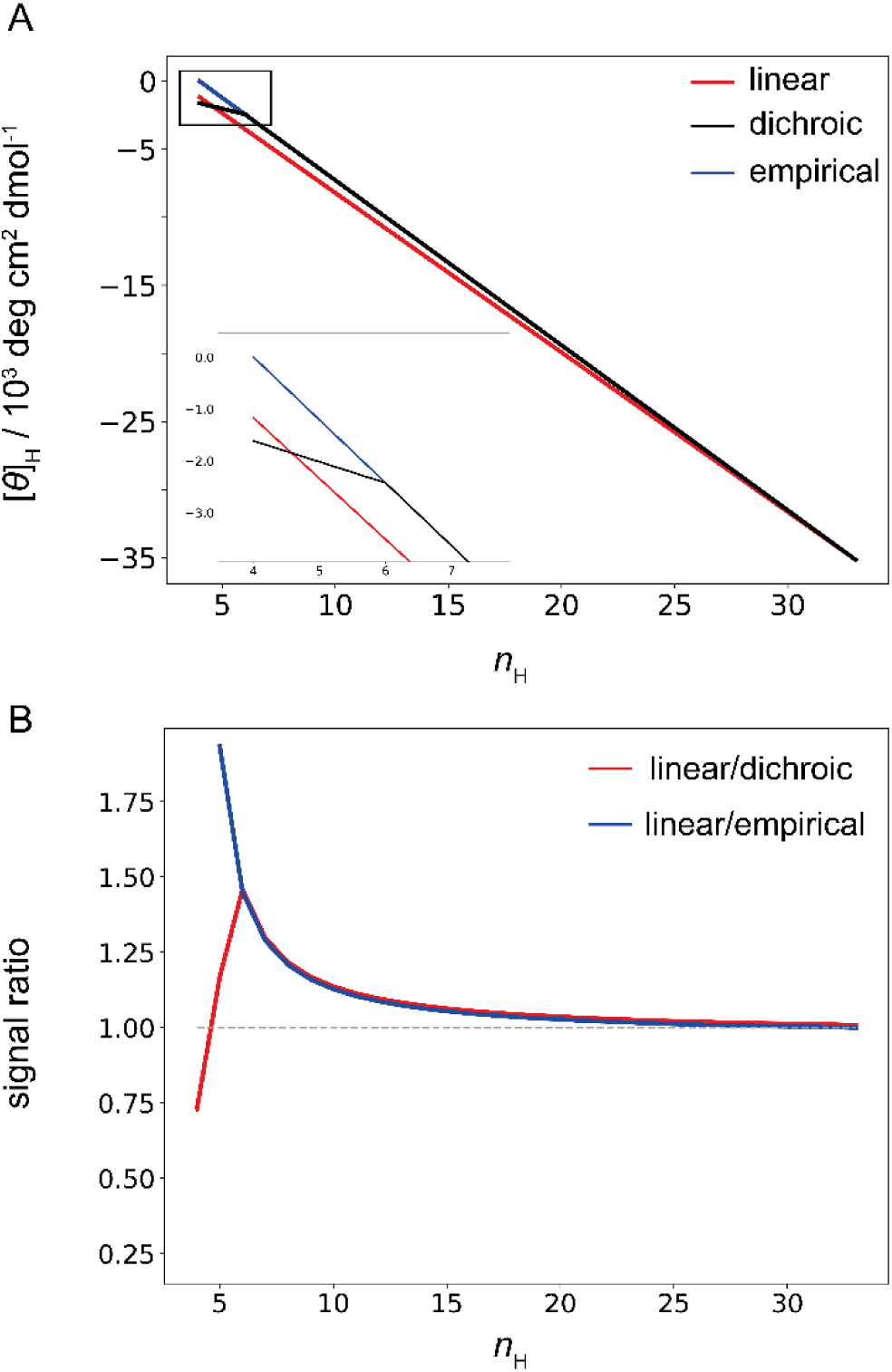
Ellipticity of peptide conformers with different lengths of helical segments. A) Helix ellipticity ([*θ*]_H_ in units per peptide bond) as a function of the number of helical units (*n*_H_) calculated with different models. Due to the constant end-effect correction approximation, the linear model fails to predict the non-linear dependence on helix length (red line) given by the two ensemble models, which evaluate the end-effect correction for each conformer individually (black and blue for dichroic and empirical, respectively). The inset shows the ellipticity of short helix segments where dichroic and empirical models diverge. B) The relative difference in the molar ellipticities calculated in the upper panel. The parameters used in calculation are the same as in Figure 1.

Dichroic and empirical models give identical [*θ*]_H_ contributions for most *n*_H_, but begin to diverge when helix segments become short *n*_H_ ≤ 6 (**Figure 2A, inset**). In these helices, all peptide units are constrained by only one hydrogen bond, so the dichroic model assigns an identical contribution [*θ*]_H1_ to all units. Essentially, in the dichroic model, the helix ellipticity becomes independent of the helix length when *n*_H_ ≤ 6. The empirical model, on the other hand, retains the length-dependence even in short helices (Eq. 2). As we show below, this is probably incorrect, as evidenced by the poor fit to the experimental data. The ratio of the CD signals estimated with the linear approximation and both ensemble models is shown in **Figure 2B** and reveals a significant relative difference for short helices (*n*_H_ = 4) and intermediate helices (4< *n*_H_ <10), for which the linear approximation leads to an underestimation of 25% and an overestimation of up to 50% of the CD signal, respectively. It should be noted that the magnitudes of all signals and consequently their differences depend on the assumed values of the end-effect parameter *k* (**Figure S1**).

Another overlooked problem of the linear model concerns peptide conformers with two or more helical segments. Although such conformers typically constitute only a minor fraction due to unfavorable helix nucleation [12], their population increases with increasing peptide length. In the linear model, double-helix conformers are spectroscopically treated as single-helix segments, although the end-effect is twice as large. Here lies the second source of signal discrepancy which again leads to overestimation of CD signal and underestimation of peptide helicity. However, in the dichroic model, spectroscopic contributions are explicitly assigned to each peptide unit, which makes accounting for multiple end-effects straightforward. We observe that for conformers with multiple helical segments, the [*θ*]_H_ values estimated with the linear model do not converge even for *n*_H,max_ (**Figure S2**). In fact, the linear model overestimates [*θ*]_H_ of double-helix conformers for the entire *n*_H_ range.

### Implementation of a matrix method for enumeration of helical conformers of varying lengths

Although the three spectroscopic models give different results for individual conformers, the relevant question is how much they differ at the level of ensemble, which consists of a mixture of conformers having different helix lengths. A traditional and well-established method for enumerating ensemble conformers is based on the matrix formalism [12]. However, the commonly used 3x3 matrix yields the joint product terms containing spectroscopically different states, which complicates the calculation of per-conformer [*θ*]_H_ contributions. As an alternative, iterative enumeration algorithms that generate every conformer one by one were previously used [9], [11]. To compare the spectroscopic models at the ensemble level, we used a modified transfer matrix that yields separate terms for the spectroscopically different states (**Methods, SI**). Briefly, instead of the conventional 3x3 matrix, we used an 8x8 matrix to separate the helical states in the coil context (*chc* and *chhc* states) from those in the helical context. The matrix product is expressed symbolically with additional statistical weights used as labeling symbols for *chc* and *chhc* states. This simplifies the classification of the different terms in the partition function polynomial and separation of the single and double helix segments. Only double-helical segments were considered, because the probability of triple or higher helical segments is negligible (< 3%) in the studied range of peptide lengths and helix-coil parameters. Collectively, the modified matrix provides a simple procedure for calculation of the length-corrected [*θ*]_H_ contributions for each peptide conformer in the ensemble.

### Ensembles with intermediate helicity and double helix segments lead to larger signal discrepancy in the linear model

Next, we examine how the spectroscopic models compare at the level of the ensemble. The probabilities of the different peptide conformers depend on the assumed helix-coil parameters *w* and *v* and the peptide length. We simulate different molecular ensembles by either changing the propagation constant *w* while keeping the nucleation constant *v* fixed, or vice versa. The propagation constant reflects how favorable is the addition of helical peptide unit to the existing helical segment. Therefore, changing *w* tunes the overall helicity of the ensemble from a predominantly coil (helix fraction ≈ 0% for *w* = 0.6) to a helical ensemble (helix content ≈ 90% for *w* = 1.8) (**Figure 3**). All three spectroscopic models predict similar [*θ*]_222_ as a function of *w* (**Figure 3A**) and converge to the identical value for the highly helical ensembles, in agreement with previous per-conformer analysis. To examine the limitations of the length-independent [*θ*]_H_ approximation used in the linear model, we calculated the signal difference between the linear and the two ensemble models. As expected, signals were overestimated in the linear model with the difference peaking for the ensembles with intermediate helix content (helix content ≈ 45%, *w ≈* 1.2) where [*θ*]_222_ differ by up to 5% (**Figure 3A, bottom**). At *w* = 1.2, the intermediate-sized helices are the dominant species (**Figure 3B**). Although the signal differences between linear and dichroic models are largest for the small helices (**Figure 2B**), these conformers make only a small absolute contribution to the [*θ*]_222_ signal. On the other hand, the longer intermediate helical segments make a larger absolute contribution to the [*θ*]_222_ signal, and although the discrepancy in signals is smaller for the intermediate helices (**Figure 2B**), the cumulative difference peaks at intermediate helicities. In other words, there is a tradeoff between relative signal difference and absolute signal intensity, which explains why the models differ most at intermediate helix contents.

**Figure 3.**
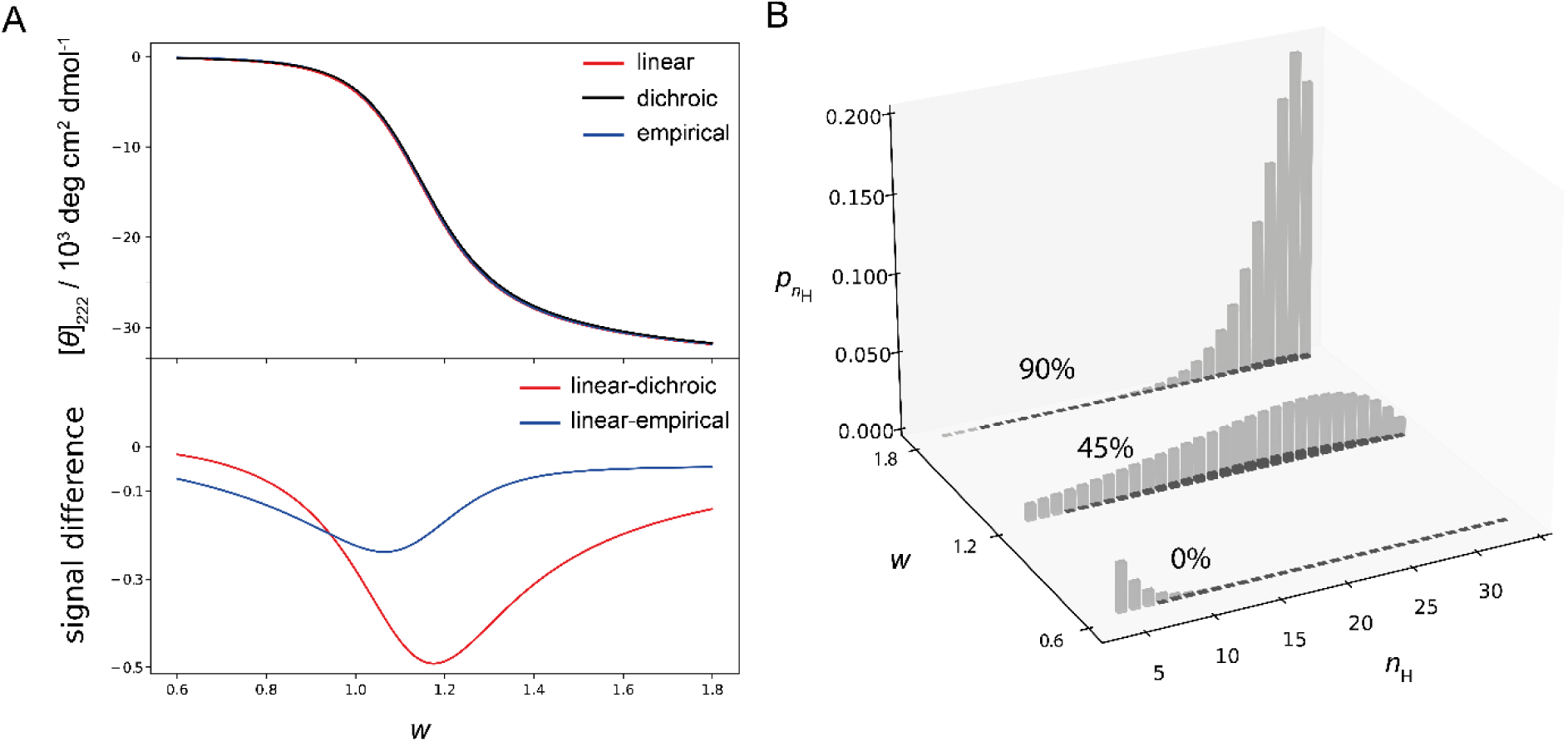
Ensemble ellipticity as a function of the propagation constant. A) Changing the propagation constant *w* tunes the overall ensemble helicity. Top panel shows the total ensemble signal calculated with different models as a function of *w* (*v* = 0.048, other parameters are the same as in Figure 1). Lower panel shows the corresponding signal difference (linear model minus dichroic or empirical). B) Ensemble composition for the selected *w* values. The gray bars show the fractional population of conformers with different number of helical units (*n*_H_). Darker bars correspond to conformers containing two helix segments. Conformers that contribute to coil (*n*_H_ < 4) are not shown.

Changes in the nucleation constant *v* can be used to analyze spectroscopic differences arising from double-helix conformers. The nucleation constant is related to the probability for initiating a helix from a segment of coil units, and the commonly used value is *v* = 0.048 [13]. As *v* changes from 0.01 to 0.05, the ensemble helicity increases and all three models give similar ellipticity (**Figure 4A**). A further increase from 0.05 to 0.1 does not increase ensemble helicity, but promotes breaking of single-helix conformers into double-helix conformers due to more favorable nucleation (**Figure 4B**). The calculated ellipticities now start to diverge (**Figure 4A, bottom**), which is due to the spectroscopic contribution of double-helix conformers. These are not specifically accounted for in linear and empirical models (the end-effect correction is applied only once). Therefore, both models overestimate the signal when ensembles contain a significant fraction of double-helix conformers (**Figure 4A**). In sum, the approximations used in the linear model lead to an overestimation of the CD signals and consequently to an underestimation of the actual helix content. This is because the linear model does not apply length-corrections to individual conformers in the ensemble and because conformers with multiple helical segments are treated as single-helix conformers. The difference in signal depends on the ensemble composition and the spectroscopic parameter *k* and appears to reach around 5% overall.

**Figure 4.**
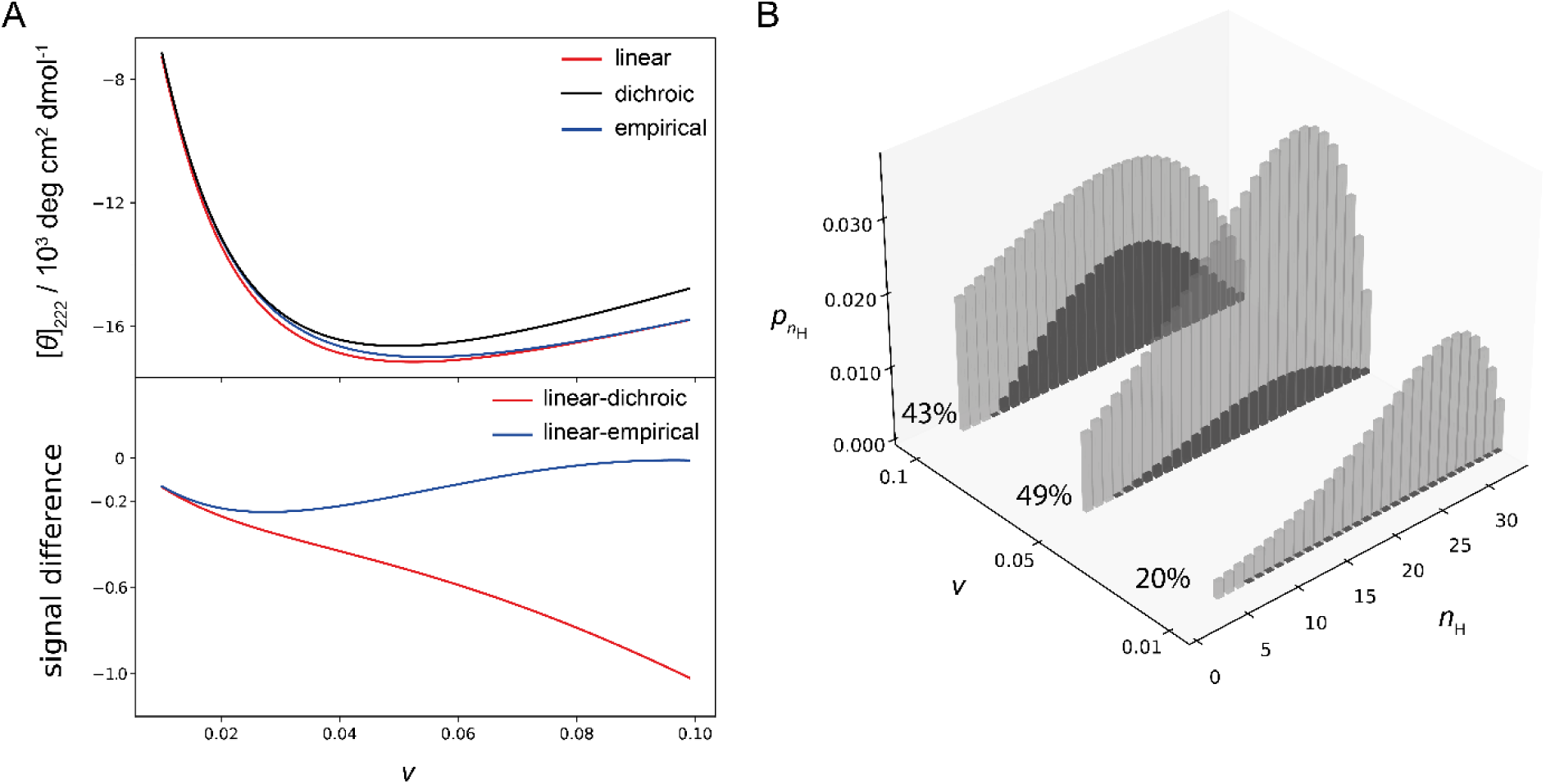
Ensemble ellipticity as a function of the nucleation constant. A) Changing the nucleation constant *v* changes the ratio between single and double-helix conformers. Top panel shows the total ensemble signal calculated with different models as a function of *v* (*w* = 1.2, other parameters are the same as in Figure 1). Lower panel shows the corresponding signal difference (linear model minus dichroic or empirical). B) Ensemble composition for the selected *v* values. The gray bars show the fractional population of conformers with different number of helical units (*n*_H_). Darker bars correspond to conformers containing two helix segments. Conformers that contribute to coil (*n*_H_ < 4) are not shown.

### Determination of helix and coil baseline parameters

Determination of helix-coil parameters from experimental CD data relies on reliable estimates of spectroscopic baseline parameters. The molar ellipticities of the coil and helix are generally considered to be temperature dependent, leading to a total of five spectroscopic parameters: [*θ*]_C_, [*θ*]_H∞_, ∂[*θ*]_H∞_/∂*T*, ∂[*θ*]_C_/∂*T*, and the end-effect parameter *k*, which we assume is temperature-independent. For the dichroic model [*θ*]_H1_ can be derived from Eq. 3 and is also temperature-dependent. These five spectroscopic parameters define the CD signal of a peptide ensemble at a given temperature. The ensemble composition is defined by four helix-coil parameters: *v*, *w*, and Δ*H* and Δ*C_p_*, which represent the change in enthalpy and heat capacity associated with helix propagation. The nucleation constant *v* is considered to be independent of temperature [14]. Evidently many parameters are required to describe the helix-coil transition monitored by CD, particularly because of the large number of spectroscopic parameters. Previously, the spectroscopic parameters were usually determined from complementary experiments or adjusted on a case-by-case basis, as discussed further below.

Here, we take a different strategy to estimate spectroscopic parameters. Rather than considering the determination of helix-coil and the spectroscopic parameters as separate problems or using some pre-determined values, we attempt to determine all parameters by global fitting of the CD dataset covering a broad range of temperatures and ensemble compositions. The major part of experimental CD data consists of the thermal melts of alanine-rich peptides with the sequence (AAKAA)_n_-GY, n=3,4,5,6. These peptides are partially helical in solution (**Figure S3**), and are an established model system for analysis of helix formation without side-chain interactions [15]. However, these data alone are not sufficient to accurately determine all parameters, as evidenced by the high parameter cross-correlations and uncertainty intervals (**Figure S4**). To constrain the parameter values, additional data that hold information about the spectroscopic behavior of fully-helical and coil-like ensembles is required. We have therefore compiled [*θ*]_222_ data from the literature for stabilized helical peptides of different lengths (**Table S3**). In these systems, such as coiled-coils and chemically stabilized helical peptides [16], all peptide units are assumed to adopt helical conformation, hence [*θ*]_222_ ≈ [*θ*]_H_(*n*_H,max_). We indeed observe the expected nonlinear dependence of [*θ*]_H_ on helix length (**Figure 5A**). The solid line shows the global fit using the dichroic model, yielding the estimates of [*θ*]_H∞_ and *k* (**Figure 5A**, results for the linear model are shown in **Figure S5** for comparison). In addition, for stable, all-helix systems with high thermal stability we determine the slope of [*θ*]_222_ with temperature (pre-transition baseline) (**Table S3**). These measurements are more conveniently expressed as the signal temperature-dependence ∂[θ]_222_/∂*T*, which in case of all-helix systems, is assumed to correspond to the helix ellipticity temperature-dependence ∂[*θ*]_H_(*n*_H,max_)/∂*T* (**Figure 5B**). Just as the ellipticity [*θ*]_H_, also ∂[*θ*]_H_(*n*_H_)/∂*T* depends on the helix length (Eq. 2) and the solid line shows the result of the global fit using the dichroic model, yielding the estimate of an infinite helix ellipticity ∂[*θ*]_H∞_/∂*T* (**Figure 5B**).

**Figure 5.**
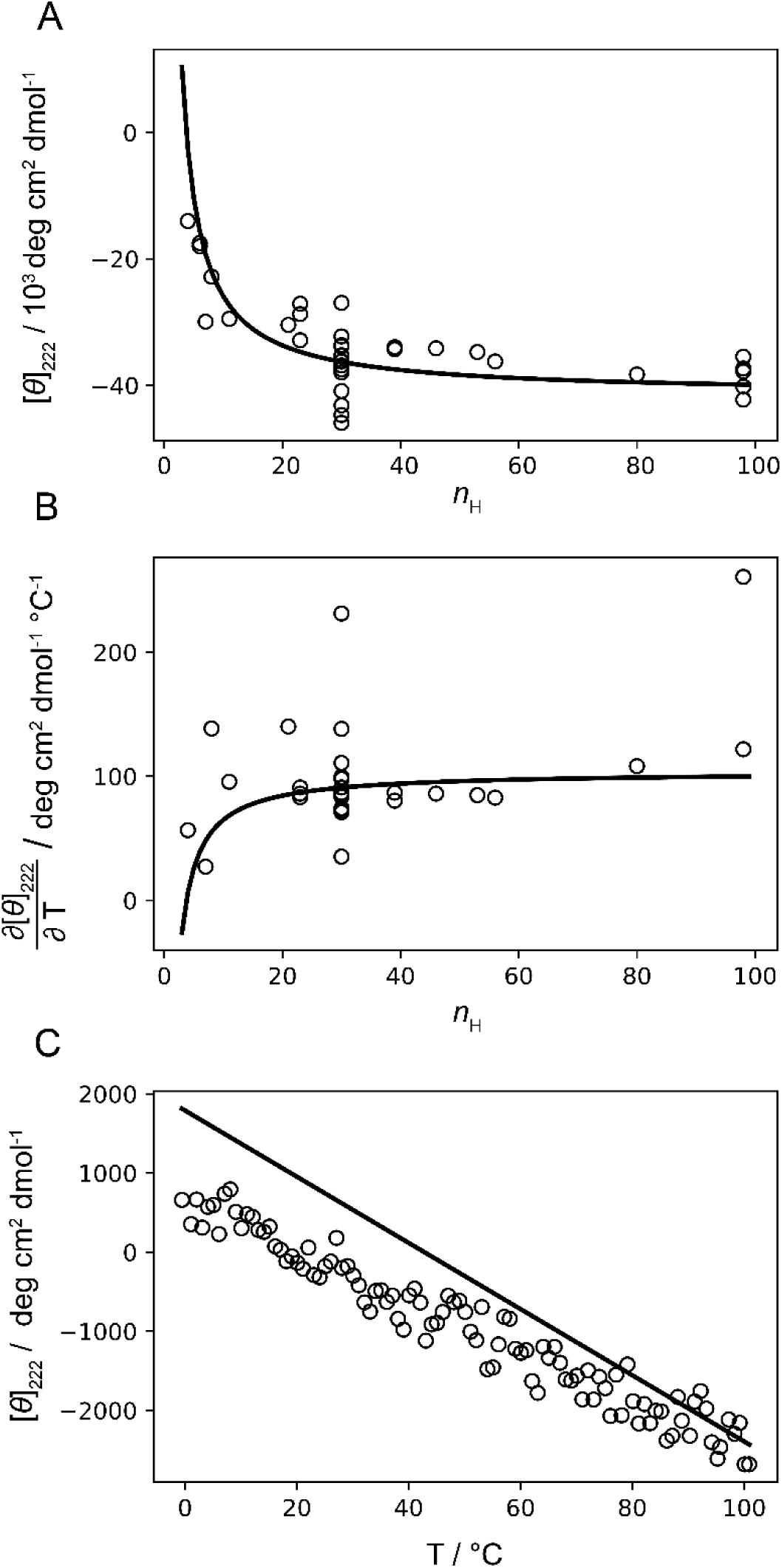
Estimation of helix and coil spectroscopic baselines. A). The measured molar ellipticity [*θ*]_222_ for 39 all-helix systems at 0°C (listed in **Table S3**) is shown as symbols. The solid line shows the helix baseline with the best-fit values [*θ*]_H∞_ and *k* obtained from global fit to all CD data using dichroic model. B) Symbols show the pretransition (*T* << *T*_m_) slopes of measured ellipticities obtained from the thermal melting curves of all-helix systems with high temperature-stability (listed in **Table S3**). The solid line shows the helix baseline temperature dependence, based on the best-fit values of ∂[*θ*]_H∞_/∂*T* and temperature-independent *k* (which also define ∂[*θ*] _H1_/∂*T*). C) Symbols show the CD thermal-melt for the AAKAA peptide, while the solid line shows the coil baseline with best-fit values [*θ*]_C_ and ∂[*θ*]_C_/∂*T*.

To constrain the coil baseline parameters, we additionally measured CD thermal melting for the short peptide (AAKAA)_n_-GY, n = 1 (abbreviated as AAKAA, **Figure 5C**). Previously, short peptides were used to determine the coil baseline, and such measurements were considered to directly correspond to the ellipticity of pure coil state ([*θ*]_222_ ≈ [*θ*]_C_). This may not be entirely correct, as even short peptides could contain helical conformers that could bias the baseline determination, especially given the large helix ellipticity. For this reason, we consider the AAKAA data in the global fit and do not impose an assumption of pure coil state for AAKAA. Indeed, the global fit using the dichroic model shows that the true coil baseline (solid line) lies above the [*θ*]_222_ data and converges to the data points at high temperatures (**Figure 5C**, results for the linear model are shown in **Figure S5** for comparison). This indeed indicates the presence of a minor fraction of helical conformers in the AAKAA ensemble at low temperatures.

### Model-analysis of helix-coil transition for alanine peptides of varying lengths

The ultimate goal of deciphering the relationship between the ensemble composition and the measured CD signal is to obtain a more precise estimate of the helix-coil parameters. We perform a global fit to all CD data simultaneously (**Figures 5 and 6**). The short-peptide and all-helix peptide dataset (**Figure 5**) mainly constrain the spectroscopic parameters, while the (AAKAA)_n_-GY n=3,4,5,6 thermal melts (**Figure 6**) hold the information about the helix-coil parameters, as these peptides sample the transition from the low-temperature helical ensembles to the high-temperature coil-like ensembles. Interestingly, linear and dichroic models give almost indistinguishable fits to the data (**Figure 6A**). Both models describe the data almost perfectly with only a small but notable deviation for the AAKAA peptide at low temperatures (**Figure 6A, inset**). The empirical model performs worse, particularly in the high-temperature region where the model-calculated ellipticity systematically deviates for all peptides (**Figure S6**). It therefore appears that the spectroscopic contribution of short helices (*n*_H_ ≤ 6), which dominate at these conditions, is not correctly predicted in the empirical model (Eq. 2).

**Figure 6.**
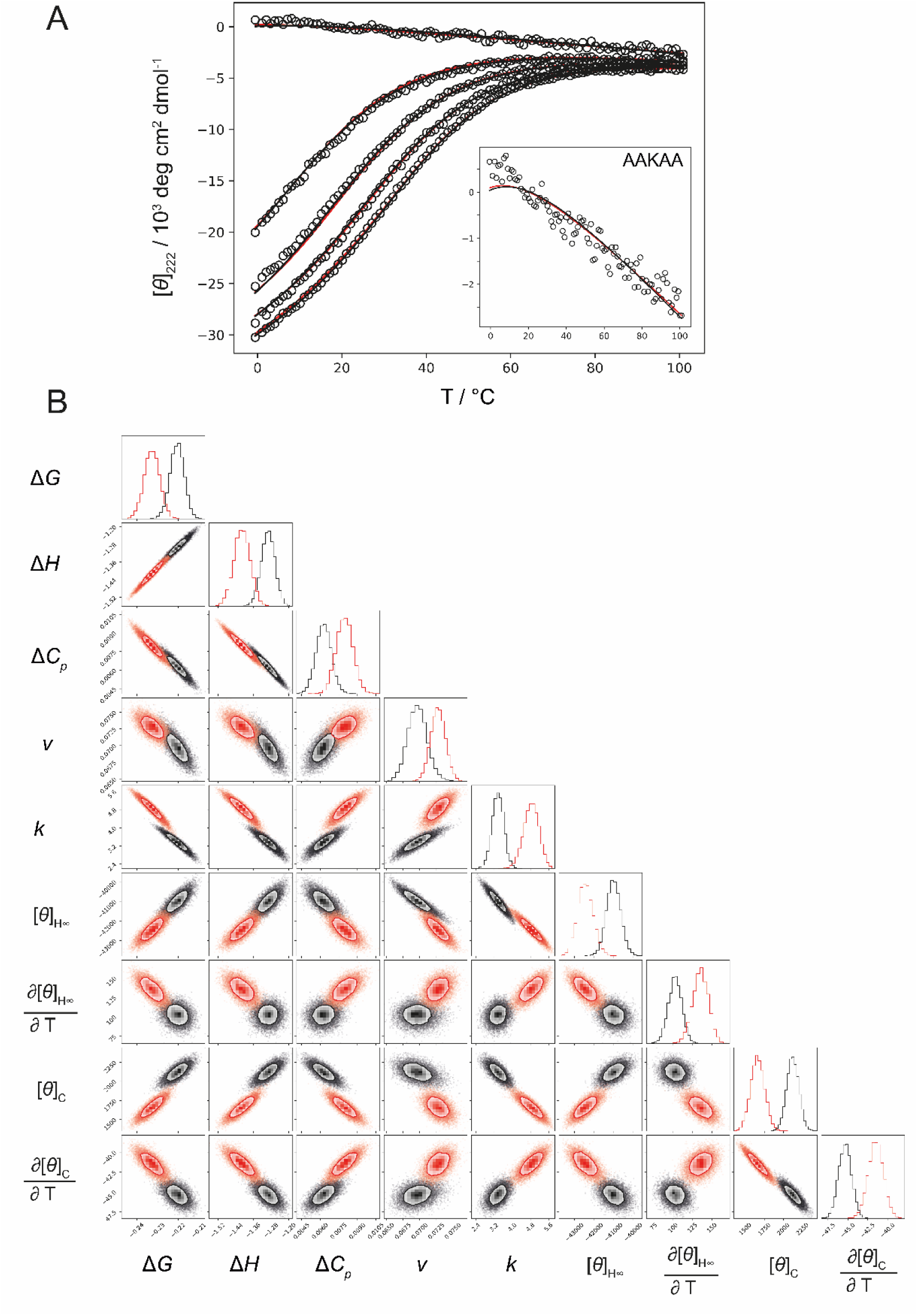
Estimation of spectroscopic and helix-coil parameters by global fitting of CD data. A) Global fit to (AAKAA)_n_-GY thermal melts with data shown as symbols. The black and red lines represent the global fit using the dichroic and linear models, respectively. The inset shows the close view of the AAKAA data and the model fits. B) MCMC corner plot for both models (linear in red and dichroic in black color) with the corresponding model parameters’ posterior distributions. The MCMC parameter statistics for the dichroic model are reported in Table 1, while for the linear model in **Table S4**.

**Table 1.** Best-fit parameters for the global fit to CD data using a dichroic spectroscopic model. Parameter values are reported per peptide unit at 0°C and correspond to the mean of MCMC-obtained posterior distributions of model parameters along with their corresponding 2σ deviations.

| Spectroscopic parameters |  | Helix-coil parameters |  |
| --- | --- | --- | --- |
| $[\theta]_{H\infty}$<br>[deg cm<sup>2</sup> dmol<sup>-1</sup>] | -41,000 ± 1,400 | $\Delta G$<br>[kcal mol<sup>-1</sup>] | -0.22 ± 0.01 |
| $\partial[\theta]_{H\infty}/\partial T$<br>[deg cm<sup>2</sup> dmol<sup>-1</sup> °C<sup>-1</sup>] | 100 ± 30 | $\Delta H$<br>[kcal mol<sup>-1</sup>] | -1.3 ± 0.1 |
| $k$ | 3.4 ± 1.0 | $\Delta C_p$<br>[kcal mol<sup>-1</sup> K<sup>-1</sup>] | 0.006 ± 0.002 |
| $[\theta]_C$<br>[deg cm<sup>2</sup> dmol<sup>-1</sup>] | 2,100 ± 300 | $\nu$ | 0.07 ± 0.01 |
| $\partial[\theta]_C/\partial T$<br>[deg cm<sup>2</sup> dmol<sup>-1</sup> °C<sup>-1</sup>] | -45 ± 5 | | |

Given the relatively worse performance of the empirical model, we focus on the fits with linear and dichroic models. We employ Bayesian inference of model parameters using Markov Chain Monte Carlo and find that despite similar quality of fit, the dichroic model provides an overall statistically better estimate of model parameters based on several metrics (**Figure 6B**). Both the uncertainty in the fitted parameters and parameter cross-correlations are lower in fits using dichroic model (**Figure S7**). Moreover, the parameter set is more robustly determined, while in the linear model parameters drift significantly upon assuming different fitting weights on the all-helix and (AAKAA)n-GY dataset (**Figure S8**). This suggests that the dichroic model can leverage small differences in the spectroscopic contributions of the conformers and thus extract more information from the experimental data. The previously discussed differences between the two models are reflected in different values of the estimated parameters. In particular, the linear model predicts higher values for the helix-coil parameters (*v, w*, Δ*H*), but this is compensated by a higher end-effect parameter that reduces the model-calculated helix ellipticity. **Table 1** reports the parameters from the global fits with the dichroic model, while those obtained with the linear models are listed in **Table S4**. Finally, we note that neither linear nor dichroic models provide a perfect description of experimental data - a systematic discrepancy for the shortest AAKAA peptide at low temperatures is noteworthy. Whether this is related to the inadequacy of the spectroscopic models or the helix-coil model remains to be investigated.

## Discussion

The main goal of this work was to establish a coherent framework for the calculation of peptide CD helicity, for the applications of helix-coil modeling and parameter determination. This entails use of an appropriate spectroscopic model and of accurate baseline parameters. We first thoroughly evaluated three spectroscopic models that relate the CD signal to the ensemble composition. To our knowledge, empirical, dichroic and linear models have not previously been compared on the same footing, making it difficult to understand which treatment of the CD signal is more appropriate. To make calculations with ensemble models more accessible, we have developed a new matrix-based method for calculation of the ensemble-weighted signal contributions that is computationally faster and easier to implement compared to the previously used enumeration algorithms [9], [11]. A version of the program that estimates peptide helix content from the measured CD ellipticity based on our implementation of dichroic models is freely available on GitHub (see Data availability). As shown here, the dichroic model gives a more accurate estimate of helix content, but requires calculation of ensemble composition using a computer program. Alternatively, the helix content can be more easily estimated using the linear model, although this will result in a slight underestimation of helix content. In this case, the measured CD signal at temperature *T* (in °C) can be converted to fractional helicity (*f*_H_) using re-calibrated spectroscopic parameters (**Table S4**) as follows:

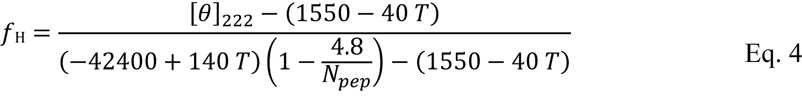

Even though the parameter values obtained from global fits using the dichroic model (**Table 1**) appear to be more accurate, when helix content is estimated using the linear model we suggest to use the corresponding set of baseline parameters (as given in Eq. 4 and Table S4).

A comparison between the linear model and the two ensemble models (dichroic, empirical) shows that the linear model tends to underestimate the true ensemble helicity (**Figures 2-4**). The difference depends on the ensemble composition and arises from the underestimation of signals of conformers with intermediate-sized helical segments and those with double-helix segments (**Figure 3 and 4**). For example, in larger peptides such as (AAKAA)_n_-GY n=6, the linear model predicts up to 5% lower helix content than the dichroic model (**Figure S9**). Nevertheless, the linear model can give an excellent fit to the data due to compensation of the model parameters (**Figure 6**). However, these parameters have higher uncertainty and cross-correlations, compared to those estimated using the dichroic model. Perhaps even more problematic aspect of parameter estimation with the linear model are the systematic drifts of parameter values depending on the assumed fitting weights on various CD datasets. This suggests that the linear model is poor at incorporating new evidence to constrain the model parameters. This leaves more room for compensation of free parameters and leads to overall parameter drifting (**Figure S7 and S8**). In contrast, for the dichroic model the parameter set appears to be stable and the cross-correlations are lower, although still high for some parameter combinations (**Figure 6**). Therefore, the dichroic model is not only more physically realistic, but it is also able to determine the model parameters more accurately overall. The ensemble-based calculation of the spectroscopic signal has the advantage of higher model sensitivity, since the subtle spectroscopic differences of the different conformers are taken into account. Interestingly, we also observed that the length-correction used in the empirical model is likely incorrect for short helical segments. While the empirical model assumes diminishing per-peptide helix ellipticity with shortening of helix segment, the dichroic model uses a constant value [*θ*]_H1_ for all segments with *n*_H_ < 6, which seems more appropriate (note that linear model also assumes constant contribution, but in the entire *n*_H_ range). Using re-calibrated values, the contribution of six peptide units at the helix ends is [*θ*]_H1_ = −17,800 deg cm^2^ dmol^-1^. Despite overall excellent fits, we still observe a systematic discrepancy between model-calculated and experimental signal for the AAKAA peptide at low temperatures (T < 20° C), suggesting that some part of the model (helix-coil or spectroscopic) involving short helix segments is still inadequate (**Figure 6A, inset**).

There is no clear consensus on the values of spectroscopic baseline parameters, although they significantly influence the estimation of helix content and helix-coil parameters. This problem appears to be somewhat obscured in the standard procedure for analysis of peptide helicity. Typically, the CD signal is first converted to fractional helicity using linear approximation and some pre-determined baseline values. Then, the fractional helicity is used as an experimental variable, not the actual CD signal. This transformation essentially eliminates the spectroscopic part of modelling, however the assumed values of baselines and, as we show here, the linear model directly influences the estimated parameters. Here we consider the estimation of the spectroscopic and helix-coil parameters as a single problem, since it is difficult to obtain ‘pure’ coil and helix states that would allow independent determination of the baseline. We have therefore assembled a diverse set of CD data and estimate both the spectroscopic and helix-coil parameters using global fitting to the raw (i.e. untransformed) CD data (**Figure 6**). Despite significant number of parameters, we show that they can be determined with sufficient reliability when supported by a diverse CD dataset (**Figure S4**). The value of the coil baseline ([*θ*]_C_= 2100 deg cm^2^ dmol^-1^, **Table 1**) is higher than that reported by Scholtz et. al. (640 deg cm^2^ dmol^-1^) [4] and is close to the value from Luo and Baldwin (2220 deg cm^2^ dmol^-1^) [17]. In these two studies, short peptides were used as a direct surrogate for the coil state. In our study we do not fix the coil baseline to the AAKAA data, rather all data support the coil determination via the global fit. We find that even in short peptides there is a small amount of helix conformers present, as the coil baseline lies above the measured ellipticities for the AAKAA peptide (**Figure 5C**).

The estimated values of the helix baseline agree well with previously published values. The re-calibrated value for the ellipticity of infinite helix is [*θ*]_H∞_ = −41,000 deg cm^2^ dmol^-1^ and that of the end-effect parameter *k* = 3.4 (**Table 1**), although the two parameters are partially correlated (**Figure 6B**). The first estimates of these parameters came from the analysis of reference protein CD spectra ([*θ*]_H∞_ = −39,500 deg cm^2^ dmol^-1^, *k* = 2-4 [7]) and theoretical studies ([*θ*]_H∞_= −45,500 deg cm^2^ dmol^-1^, *k* = 2), while more recent theoretical studies suggest [*θ*]*_H∞_* = −37.000, and *k* = 2.6-3 [18]. Stabilization of the helical conformation using co-solvents for peptides of different lengths gave [*θ*]_H∞_ = −41,500 and *k* = 3.7, which is in excellent agreement with our estimate [9]. Studies which fit helix-coil models to synthetic peptides often use [*θ*]_H∞_ = −42,500 and *k* = 2.5. Higher values [*θ*]_H∞_ = −44,000 deg cm^2^ dmol^-1^ and *k* = 3 have been proposed by Luo and Baldwin [17], who also revised the ∂[*θ*]_H∞_/∂*T* value from previous +100 to +250 deg cm^2^ dmol^-1^ °C^-1^. However, these studies neglected the length-dependence of the helix ellipticity temperature derivate (**Figure 5B**). Unfortunately, the assembled ∂[*θ*]_222_/∂*T* data show considerable scatter obscuring the detailed nature of the helix ellipticity temperature-dependence. We therefore considered *k* as temperature-independent parameter and obtained ∂[*θ*]_H∞_/∂*T* = 100 and ∂[*θ*]_H1_/∂*T* = 40 deg cm^2^ dmol^-1^ (**Table 1**). When *k* is assumed as temperature-independent, this means that the ellipticities of single and double-bonded helical peptides change synchronously with temperature, such that their ratio (Eq. 3) stays constant at different temperatures, consequently also *k* remains unchanged. However, it is quite possible that the ellipticity ratio in Eq. 3 changes with temperature, making *k* temperature-dependent parameter. Overall, helix spectroscopic values reported here agree with previous estimates, although there is room for improvement, particularly with respect to the temperature-dependence of the helix baseline.

**Table 1** lists the helix-coil parameters for alanine-rich peptides, which provide an excellent model system for studying the thermodynamic forces behind the backbone-driven helix folding (without sidechain interactions). The reported value of helix propagation and its enthalpy contribution *w* = 1.50 and Δ*H* = −1.3 kcal mol^-1^ at 0° C agree well with previous estimates for alanine peptides [4], [17], [19]. Of particular interest are the estimates of two parameters listed in **Table 1**: Δ*C_p_* and *v.* The first is the change in heat capacity upon helix folding, a parameter that has eluded precise determination. We estimate Δ*C_p_* = +6 cal mol^-1^ K^-1^. Although it shows a significant cross-correlation with some parameters (**Figure 6B**), the fits are worse when Δ*C_p_* = 0 is assumed (**Figure S10**), supporting the existence of a small, non-negative Δ*C_p_*. Previous calorimetric studies using DSC and ITC were unable to determine the exact value of Δ*C_p_* and concluded that Δ*C_p_* is zero or very small [20], [21]. Two other studies reported a small negative heat capacity for helix folding around −4 cal^-1^ mol K^-1^ res^-1^ [22], [23]. In a study using peptides with XEARA sequence, the calorimetric Δ*H* was found to be temperature-dependent, and Δ*C_p_* = +8 cal mol^-1^ K^-1^ was estimated [24]. This value is similar to ours, although the peptide sequences differ. This perhaps suggests that Δ*C_p_* arises only from backbone interactions, but it remains to be investigated what is the molecular origin of the heat capacity increment in helix folding. The second value of interest is that of the nucleation constant *v* = 0.07, which is substantially higher than previous estimates (*v* = 0.048 [13]). Our estimate appears to be reliable in terms of low cross-correlations and is also supported by a poor fit quality when the usual *v* = 0.048 is assumed (**Figure S11**). In most helix-coil studies, the nucleation constant is used as a fixed parameter, *v* = 0.048 for the homopolymer model or *v* = 0.036 in helix-coil models with capping interactions [25]. These values were obtained from amide proton exchange experiments [13]. Our value is closer that reported by Scholtz obtained as a fitting parameter of the CD thermal melts [4] and to a more recent analysis using the helix-coil model with added solvent-effect [26]. Overall, the traditionally used value *v* = 0.048 appears to be underestimated, suggesting that the change in backbone conformational entropy is smaller than anticipated previously. This could have important implications for protein folding, as the decrease in conformational entropy is considered to be the single largest force opposing protein folding [27].

## Materials and Methods

### Peptides and sample preparation

The peptides were purchased from ChinaPeptides Ltd. and were at least 95% pure. All contained N-terminal acetyl and C-terminal amide modifications. Peptides were dissolved in mili-Q water at about 1-3-mg/ml concentration and dialyzed against 10mM phosphate pH 7.0, 1M NaCl buffer (phosphate buffer). Before each experiment these stock solutions were centrifuged and the peptide concentration was determined by measuring absorbance at 275 nm, using extinction coefficient 1450 M^-1^ cm^-1^ [28]. Samples were prepared by diluting stock solutions to a concentration of 100 µM, which were used in the CD experiments.

### Circular dichroism

Circular dichroism measurements were performed using a Jasco J-1500 CD spectrophotometer. All the samples were measured in 1 mm quartz cuvette (Hellma) in phosphate buffer. Ammonium d-camphor-10-sulfonate (Jasco Ltd.) was measured before and after each measurement to calibrate the intensity of the Xe-ultraviolet lamp. A blank measurement (buffer in quartz cuvette) was subtracted from each measurement. For thermal melts, ellipticity at 222 nm was measured every 1 °C with an integration time of 8 seconds. Reversibility was checked by comparing spectra before and after denaturation and all samples showed complete signal reversibility. The raw signal (ellipticity) was converted to mean molar ellipticity [*θ*]_222_ of peptide unit, according to the following equation:

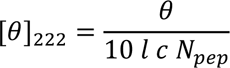

where *θ* is the measured ellipticity (raw signal) in mdeg, *l* is the optical length in cm, *c* is the peptide molar concentration, and *N*_pep_ is the number of peptide units in peptide, which in case of N- and C-terminally modified peptides equals *N*_pep_ = *N*_res_ + 1.

### All-helix CD dataset

The literature was searched for suitable coiled-coil systems or short stabilized peptides that remain fully helical over a wide temperature range (high melting temperature). Thermal melting curves were digitized using computer program PlotDigitizer to extract the values [*θ*]_222,0°C_ and ∂[*θ*]_222_ /∂*T* [29]. Details of the data points in the dataset are summarized in **Table S1**.

### Lifson-Roig model

In the LR model, each residue in the peptide chain can be either helical (*h*) or coil (*c*) based on the geometry of the backbone (Φ, Ψ angles). This results in 2*^N^* states for a peptide with *N* units. All of these conformers are enumerated using the matrix method, resulting in an analytical expression for the partition function *Q*. Two parameters determine the partition function: the nucleation parameter *v* and the propagation parameter *w*. The nucleation parameter *v* is assumed to be constant, mainly entropic by origin, and temperature-independent, while the helix propagation parameter *w* is related to the formation of backbone hydrogen bond and is considered to be temperature-dependent [12]. The probability for the *i*-th conformer can be calculated by dividing its statistical weight with the partition function of the whole peptide ensemble.

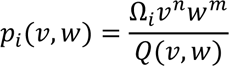

The conformer contains *n* and *m* units having weights *v* and *w*, while Ω_i_ is the total number of possible states with the same composition (degeneracy). Given the probability for each conformer in the ensemble, the CD signal can be calculated using Eq. 1. More details on the model implementation can be found in the Supplementary information. All calculations were performed using custom Python scripts relaying heavily on numpy and sympy library [30], [31].

### Global fitting and Statistical analysis

A Bayesian inference analysis of the models was performed using Markov Chain Monte Carlo (MCMC). The analysis was performed using Python probabilistic programming library PyMC [32]. We adopt a statistical model for the CD data as being generated by general model function presented in Eq. 1 with added Gaussian noise (𝜀).

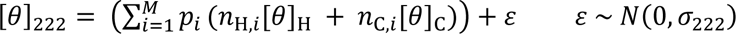

where 𝜎_222_ expresses experimental scatter of the measured and acquired CD data. For all the model parameters we set non-informative (uniform) priors, that allow model parameters to explore entire parameter space during MCMC without constrains.

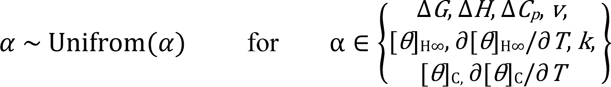

NUTS (No-U-Turn) sampler was used to infer posterior distributions of model parameters with 3000 samples (1000 burn-in). Correlation coefficients were calculated as Spearman’s rank correlation coefficients.

### Data availability

All data are included in the paper and Supplementary information. A computer code that implements dichroic model for calculation of peptide helix content based on the measured CD ellipticity is available on GitHub at https://github.com/sanhadzi/Dichroic-CD-model.

## Acknowledgments

This work was supported by the Grants P1-0201 and J1-50026 from the Slovenian Research Agency to S.H. and J.L. U.Z. acknowledges support through the “Young Researchers” program of the Slovenian Research Agency.

## References

[1] C. Toniolo, F. Formaggio, and R. W. Woody, “Electronic Circular Dichroism of Peptides,” Compr. Chiroptical Spectrosc. Appl. Stereochem. Anal. Synth. Compd. Nat. Prod. Biomol., pp. 499–544, Jan. 2012.

[2] D. M. Rogers, S. B. Jasim, N. T. Dyer, F. Auvray, M. Réfrégiers, and J. D. Hirst, “Electronic Circular Dichroism Spectroscopy of Proteins,” Chem, vol. 5, no. 11, pp. 2751–2774, Nov. 2019.

[3] R. W. Woody, “[4] Circular dichroism,” Methods Enzymol., vol. 246, no. C, pp. 34–71, Jan. 1995.

[4] J. M. Scholtz, H. Qian, E. J. York, J. M. Stewart, and R. L. Baldwin, “Parameters of helix-coil transition theory for alanine-based peptides of varying chain lengths in water,” Biopolymers, vol. 31, no. 13, pp. 1463–1470, Nov. 1991.

[5] P. Luo and R. L. Baldwin, “Mechanism of helix induction by trifluoroethanol: a framework for extrapolating the helix-forming properties of peptides from trifluoroethanol/water mixtures back to water,” Biochemistry, vol. 36, no. 27, pp. 8413–8421, Jul. 1997.

[6] N. R. Kallenbach, P. Lyu, and H. Zhou, “CD Spectroscopy and the Helix-Coil Transition in Peptides and Polypeptides,” Circ. Dichroism Conform. Anal. Biomol., pp. 201–259, 1996.

[7] C. YH, Y. JT, and C. KH, “Determination of the helix and beta form of proteins in aqueous solution by circular dichroism,” Biochemistry, vol. 13, no. 16, pp. 3350–3359, Jul. 1974.

[8] M. C. Manning and R. W. Woody, “Theoretical CD studies of polypeptide helices: examination of important electronic and geometric factors,” Biopolymers, vol. 31, no. 5, pp. 569–586, 1991.

[9] P. J. Gans, P. C. Lyu, M. C. Manning, R. W. Woody, and N. R. Kallenbach, “The helix-coil transition in heterogeneous peptides with specific side-chain interactions: theory and comparison with CD spectral data,” Biopolymers, vol. 31, no. 13, pp. 1605– 1614, 1991.

[10] W. Kauzmann and H. Eyring, “The Effect of the Rotation of Groups about Bonds on Optical Rotatory Power,” J. Chem. Phys., vol. 9, no. 1, pp. 41–53, Jan. 1941.

[11] W. Shalongo and E. Stellwagen, “Dichroic Statistical Model for Prediction and Analysis of Peptide Helicity,” Proteins, vol. 28, pp. 467–480, 1997.

[12] D. (Douglas) Poland and H. A. Scheraga, Theory of helix-coil transitions in biopolymers; statistical mechanical theory of order-disorder transitions in biological macromolecules,. Academic Press, 1970.

[13] C. A. Röhl, J. Martin Scholtz, R. L. Bald, E. J. York, and J. M. Stewart, “Kinetics of amide proton exchange in helical peptides of varying chain lengths. Interpretation by the Lifson-Roig Equation,” Biochemistry, vol. 31, no. 5, pp. 1263–1269, Feb. 1992.

[14] H. Qian and J. A. Schell, “Helix-Coil Theories: A Comparative Study for Finite Length Polypeptides,” Phys. Chem, vol. 96, pp. 3987–3994, 1992.

[15] S. Marqusee, V. H. Robbins, and R. L. Baldwin, “Unusually stable helix formation in short alanine-based peptides.,” Proc. Natl. Acad. Sci. U. S. A., vol. 86, no. 14, pp. 5286–5290, Jul. 1989.

[16] N. E. Shepherd, H. N. Hoang, G. Abbenante, and D. P. Fairlie, “Single turn peptide alpha helices with exceptional stability in water,” J. Am. Chem. Soc., vol. 127, no. 9, pp. 2974–2983, Mar. 2005.

[17] P. Luo and R. L. Baldwin, “Mechanism of helix induction by trifluoroethanol: A framework for extrapolating the helix-forming properties of peptides from trifluoroethanol/water mixtures back to water,” Biochemistry, vol. 36, no. 27, pp. 8413–8421, Jul. 1997.

[18] N. A. Besley and J. D. Hirst, “Theoretical studies toward quantitative protein circular dichroism calculations,” J. Am. Chem. Soc., vol. 121, no. 41, pp. 9636–9644, Oct. 1999.

[19] ‡ Carol A. Rohl* and R. L. Baldwin, “Comparison of NH Exchange and Circular Dichroism as Techniques for Measuring the Parameters of the Helix−Coil Transition in Peptides†,” 1997.

[20] J. M. Scholtz et al., “Calorimetric determination of the enthalpy change for the alpha-helix to coil transition of an alanine peptide in water.,” Proc. Natl. Acad. Sci., vol. 88, no. 7, pp. 2854–2858, Apr. 1991.

[21] M. M. Lopez, D. H. Chin, R. L. Baldwin, and G. I. Makhatadze, “The enthalpy of the alanine peptide helix measured by isothermal titration calorimetry using metal-binding to induce helix formation,” Proc. Natl. Acad. Sci. U. S. A., vol. 99, no. 3, pp. 1298– 1302, Feb. 2002.

[22] J. W. Taylor, N. J. Greenfield, B. Wu, and P. L. Privalov, “A calorimetric study of the folding-unfolding of an alpha-helix with covalently closed N and C-terminal loops,” J. Mol. Biol., vol. 291, no. 4, pp. 965–976, Aug. 1999.

[23] W. Shalongo, L. Dugad, and E. Stellwagen, “Analysis of the Thermal Transitions of a Model Helical Peptide Using 13C NMR,” J. Am. Chem. Soc., vol. 116, no. 6, pp. 2500–2507, Mar. 1994.

[24] J. M. Richardson and G. I. Makhatadze, “Temperature Dependence of the Thermodynamics of Helix–Coil Transition,” J. Mol. Biol., vol. 335, no. 4, pp. 1029– 1037, Jan. 2004.

[25] C. A. Rohl and R. L. Baldwin, “Deciphering rules of helix stability in peptides,” Methods Enzymol., vol. 295, pp. 1–26, 1998.

[26] K. Yeritsyan, M. Valant, and A. Badasyan, “Processing helix–coil transition data: Account of chain length and solvent effects,” Front. Nanotechnol., vol. 4, p. 73, Oct. 2022.

[27] K. A. Dill, “Dominant forces in protein folding,” Biochemistry, vol. 29, no. 31, pp. 7133–7155, Aug. 1990.

[28] J. F. Brandts and L. J. Kaplan, “Derivative Spectroscopy Applied to Tyrosyl Chromophores. Studies on Ribonuclease, Lima Bean Inhibitors, Insulin, and Pancreatic Trypsin Inhibitor,” Biochemistry, vol. 12, no. 10, pp. 2011–2024, May 1973.

[29] A. Rohatgi, “Webplotdigitizer: Version 4.6,” 2022. [Online]. Available: https://automeris.io/WebPlotDigitizer.

[30] C. R. Harris et al., “Array programming with NumPy,” Nature, vol. 585, no. 7825. Nature Research, pp. 357–362, 17-Sep-2020.

[31] A. Meurer et al., “SymPy: symbolic computing in Python,” PeerJ Comput. Sci., vol. 3, p. e103, Jan. 2017.

[32] O. Abril-Pla et al., “PyMC: a modern, and comprehensive probabilistic programming framework in Python,” PeerJ Comput. Sci., vol. 9, p. e1516, Sep. 2023.

