## Supplementary information for "Analysis of the peptide helicity using an ensemble spectroscopic model with re-calibrated parameters"

for

### Implementation of dichroic spectroscopic model

To construct the partition functions of the Lifson-Roig (LR) model for peptides, we used a larger 8x8 LR transfer matrix (*M*) [1]. This matrix relates four adjacent residues and gives statistical weights for different conformers of the quadruplet (**Table S1**).

$$M = \begin{bmatrix} w & v & 0 & 0 & 0 & 0 & 0 & 0 \\ 0 & 0 & 1 & 1 & 0 & 0 & 0 & 0 \\ 0 & 0 & 0 & 0 & v & X & 0 & 0 \\ 0 & 0 & 0 & 0 & 0 & 0 & 1 & 1 \\ w & Y & 0 & 0 & 0 & 0 & 0 & 0 \\ 0 & 0 & 1 & 1 & 0 & 0 & 0 & 0 \\ 0 & 0 & 0 & 0 & v & X & 0 & 0 \\ 0 & 0 & 0 & 0 & 0 & 0 & 1 & 1 \end{bmatrix}$$

Two additional weights *X* and *Y* in the matrix *M* denote the states in which the *i*-th residue is in the helical conformation, surrounded by residues in the coil conformation. **Table S1** lists all 16 possible combinations for four adjacent residues with the corresponding statistical weights.

**Table S1**. Assignments of the 16 possible states of four neighboring residues. Statistical weights are assigned to each of the states. Note that states with single *h* and *hh* are given special labeling weights *X* and *Y*, respectively.

| States | | | | |
|---|---|---|---|---|
| ***i*-2** | ***i*-1** | ***i*** | ***i*+1** | **Statistical weight of the *i*<sup></sup>th residue** |
| *h* | *h* | *h* | *h* | w |
| *h* | *h* | *h* | *c* | v |
| *h* | *h* | *c* | *h* | 1 |
| *h* | *h* | *c* | *c* | 1 |
| *h* | *c* | *h* | *h* | v |
| *h* | *c* | *h* | *c* | X |
| *h* | *c* | *c* | *h* | 1 |
| *h* | *c* | *c* | *c* | 1 |
| *c* | *h* | *h* | *h* | w |
| *c* | *h* | *h* | *c* | Y |
| *c* | *h* | *c* | *h* | 1 |
| *c* | *h* | *c* | *c* | 1 |
| *c* | *c* | *h* | *h* | v |
| *c* | *c* | *h* | *c* | X |
| *c* | *c* | *c* | *h* | 1 |
| *c* | *c* | *c* | *c* | 1 |

Partition function *Q* for *N*-residue long peptide is then calculated via standard matrix-multiplication procedure.

$$Q = m^{+}\left(\prod_{i=1}^{N} M_i\right) m^{-} = m^{+} M^{\mathrm{N}} m^{-}$$

where $m^{+} = (0,0,0,0,0,0,1,1)$ and $m^{-} = (0,1,0,1,0,1,0,1)^{T}$ are the end vectors, that ensure that last units in the chain (N- and C- terminal groups) can only be in the coil state (weight 1). After expanding matrix product, we obtain partition function polynomial. Below is an example for chain with ten units ($N = 10$).

$Q$ ($N = 10$) =
6*X**5 + 5*X**4*Y*v + 35*X**4 + 60*X**3*Y*v + 4*X**3*v**2*w**2 + 20*X**3*v**2*w + 56*X**3 + 30*X**2*Y**2*v**2 + 12*X**2*Y*v**3*w + 105*X**2*Y*v + 3*X**2*v**2*w**4 + 12*X**2*v**2*w**3 + 30*X**2*v**2*w**2 + 60*X**2*v**2*w + 36*X**2 + 4*X*Y**3*v**3 + 60*X*Y**2*v**2 + 6*X*Y*v**3*w**3 + 24*X*Y*v**3*w**2 + 60*X*Y*v**3*w + 56*X*Y*v + 6*X*v**4*w**3 + 12*X*v**4*w**2 + 2*X*v**2*w**6 + 6*X*v**2*w**5 + 12*X*v**2*w**4 + 20*X*v**2*w**3 + 30*X*v**2*w**2 + 42*X*v**2*w + 10*X + 10*Y**3*v**3 + 3*Y**2*v**4*w**2 + 12*Y**2*v**4*w + 21*Y**2*v**2 + 3*Y*v**5*w**2 + 2*Y*v**3*w**5 + 6*Y*v**3*w**4 + 12*Y*v**3*w**3 + 20*Y*v**3*w**2 + 30*Y*v**3*w + 9*Y*v + 4*v**4*w**5 + 9*v**4*w**4 + 12*v**4*w**3 + 10*v**4*w**2 + v**2*w**8 + 2*v**2*w**7 + 3*v**2*w**6 + 4*v**2*w**5 + 5*v**2*w**4 + 6*v**2*w**3 + 7*v**2*w**2 + 8*v**2*w + 1

From such a partition function polynomial, we can easily identify spectroscopically relevant states – e.g., single, double helices, since the weight *v* is now present only in polynomial terms that have end of the helices or doublet *hh* states. Note however, that for the case of *chhc*, two statistical weights are assigned (*Y*v*). Therefore, the polynomial term 2**Y*v***3**w***5 for example stands for a single helix *v***2**w***5 with an additional isolated helix "doublet" (*hh*) in the chain. Integer numbers represent the degeneracy of such a conformer in the partition function. As mentioned earlier, the *X,Y* weights were used for labeling purposes only and otherwise considered equal to *v* in all calculations (*X, Y* = *v*).

### Ellipticity contributions

Given the partition function polynomial *Q*, implementation of single helical contribution to the CD signal is straightforward. Here is the general scheme for single helix CD contribution.

$$[\theta]_{single\ helix}\,(n_{\mathrm{w}}) = \begin{cases} (n_{\mathrm{w}} + 3)[\theta]_{\mathrm{H1}}, & n_{\mathrm{w}} < 4 \\ 6[\theta]_{\mathrm{H1}} + (n_{\mathrm{w}} - 3)[\theta]_{\mathrm{H\infty}}, & n_{\mathrm{w}} \geq 4 \end{cases}$$

Where $[\theta]_{\mathrm{H\infty}}$ and $[\theta]_{\mathrm{H1}}$ are specific molar residual ellipticities for double- and single- H-bonded peptide units and are related by Eq. 3 in the main text. $n_{\mathrm{w}}$ is the number of *w*-weighted residues (power over *w* weight in the polynomial term). Calculation of end-corrected single helix ellipticity using the empirical model proceeds by substituting $n_{\mathrm{w}}$ into Eq. 2 for each conformer.

Calculation of the CD contribution for the double helix is a bit more complicated. Based on the composition of the individual terms in the partition function polynomial, we can easily assign the helix multiplicity of the individual terms and the total helical content. The composition (number of *w,v,X,Y* elements) of the individual terms does not directly tell us anything about the individual lengths of the two helices in the conformer. Only the total of two helix segments is known. For example, the term $v^4w^6$ contains all the combinations of double helices that add to the sixth power of $w$ – 1:5, 2:4 and 3:3. Although different length combinations have the same probability (determined only by $v$ and $w$), they contribute to CD signal differently. However, since each combination is equally likely, we can calculate the average CD contribution of all combinations. Below is an example table with calculations of the spectroscopic contribution for a term with two helices with a total of 13 $w$-weighted residues ($v^4w^{13}$).

**Table S2.** Assignations of the CD contribution for double helix term ($v^4w^{13}$). $n_{H1}$ and $n_{H2}$ represent the number of single- and double bonded helical peptide units that contribute $[\theta]_{H1}$ and $[\theta]_{H\infty}$, respectively.

| helix-length combination | | $n_{H1}$ | $n_{H2}$ | $n_{H1} + n_{H2}$ | $<n_{H1}> + <n_{H2}>$ | **overall CD contribution $<[\theta]_{term}>$** |
|---|---|---|---|---|---|---|
| **$v^2w$** | **$v^2w^{12}$** | 4+6 | 0+9 | 10 + 9 | $11.5\ n_{H1} + 7.5\ n_{H2}$ | $P_i\ (11.5\ [\theta]_{H1} + 7.5\ [\theta]_{H\infty})$ |
| **$v^2w^2$** | **$v^2w^{11}$** | 5+6 | 0+8 | 11 + 8 | | |
| **$v^2w^3$** | **$v^2w^{10}$** | 6+6 | 0+7 | 12 + 7 | | |
| **$v^2w^4$** | **$v^2w^9$** | 6+6 | 1+6 | 12 + 7 | | |
| **$v^2w^5$** | **$v^2w^8$** | 6+6 | 2+5 | 12 + 7 | | |
| **$v^2w^6$** | **$v^2w^7$** | 6+6 | 3+4 | 12 + 7 | | |

Since the probability is the same for each substate of helix-length combination, we can average the contributions from CD across all substate combinations and get $<[\theta]_{term}>$. With the total probability ($P_i$) of the polynomial term and the averaged $<[\theta]_{term}>$, we can now perform the calculations of the sum according to Eq. 1 in the main text.

It turns out that CD contribution of each polynomial term for the double helix (as shown in **Table S2**) can be summarized by two rules used to calculate the average number of single ($<n_{H1}>$) and double ($<n_{H2}>$) H-bonded peptide units.

$$\langle n_{\mathrm{H1}} \rangle = \frac{12\ (n_{combos} - 2) + 11 + 10}{n_{combos}} \qquad \text{Eq. S1}$$

$$\langle n_{\mathrm{H2}} \rangle = \frac{(n_w - 6)(n_{combos} - 2) + (n_w - 5) + (n_w - 4)}{n_{combos}} \qquad \text{Eq. S2}$$

where $n_{w}$ is the power over $w$ in a given polynomial term and $n_{combos}$ is the number of different combinations of double helices that add up to the total length of the term. For previous example presented in **Table S2** ($v^4w^{13}$) $n_{combos}$ is 6 (helix lengths combinations 1:12, 2:11, 3:10, 4:9, 5:8, 6:7).

Equipped with all this we can write a general calculation scheme for calculation of CD contribution of double helices.

$$[\theta]_{double\ helix}\,(n_{\mathrm{w}}) = \begin{cases} (n_w + 6)[\theta]_{\mathrm{H1}}, & n_{\mathrm{w}} < 5 \\ 10.5\,[\theta]_{\mathrm{H1}} + 0.5\,[\theta]_{\mathrm{H\infty}}, & n_{\mathrm{w}} = 5 \\ \langle n_{\mathrm{H1}}\rangle[\theta]_{\mathrm{H1}} + \langle n_{\mathrm{H2}}\rangle[\theta]_{\mathrm{H\infty}}, & n_{\mathrm{w}} > 5 \end{cases}$$

where $n_{w}$ is the power over $w$ in polynomial term (number of $w$-weighted residues). $<n_{H1}>$ and $<n_{H2}>$ are the average number of single- and double- H-bonded residues that contribute $[\theta]_{H1}$ and $[\theta]_{H\infty}$ to the signal and are calculated according to the equations Eq. S1 and Eq. S2, respectively.

Calculation can be in similar manner extended to n-multiple helices, however we find only double helices to be relevant for our range of chain lengths (up to N $\approx$ 50) and helix-coil parameters.

**Table S3.** Dataset of stabilized all-helix peptides from literature and corresponding values of $[\theta]_{222}$ and $\partial[\theta]_{222}/\partial T$ obtained from the published data. Data are given per mole of peptide unit and values of $[\theta]_{222}$ are at 0 °C.

| system | $[\theta]_{222}$ [$10^3$ deg cm$^2$ dmol$^{-1}$] | $\partial[\theta]_{222}/\partial T$ [$10^3$ deg cm$^2$ dmol$^{-1}$] | length (residues) | Source (DOI) | |
|---|---|---|---|---|---|
| P3P4_2xSHn | -44.7 | 0.23 | 30 | 10.1021/jacs.7b01690 | [2] |
| P3P4_2xSHc | -45.9 | 0.23 | 30 | 10.1021/jacs.7b01690 | [2] |
| P5P6_SHcSHn | -43.2 | 0.24 | 30 | 10.1021/jacs.7b01690 | [2] |
| P5P6_2xSHn | -40.9 | 0.25 | 30 | 10.1021/jacs.7b01690 | [2] |
| EK3 | -37.3 | 0.42 | 98 | 10.1038/srep44341 | [3] |
| EK2R2 | -42.3 | 0.12 | 98 | 10.1038/srep44341 | [3] |
| EK2R1 | -40.2 | 0.26 | 98 | 10.1038/srep44341 | [3] |
| M6R | -37.8 | 0.27 | 98 | 10.1038/srep44341 | [3] |
| M6WT | -35.5 | 0.30 | 98 | 10.1038/srep44341 | [3] |
| apCC_Tet | -35.2 | 0.07 | 30 | 10.1021/jacs.8b13354 | [4] |
| CC-Hex*-II | -27.0 | 0.04 | 30 | 10.1021/jacs.8b13354 | [4] |
| CC-Hex*-L24Nle | -36.2 | 0.14 | 30 | 10.1021/jacs.8b13354 | [4] |
| cc_tet | -34.0 | 0.09 | 39 | 10.1038/nchembio.692 | [5] |
| 1EK4 | -32.3 | 0.10 | 30 | 10.1021/acs.biochem.0c00082 | [6] |
| 1KE4 | -38.0 | 0.09 | 30 | 10.1021/acs.biochem.0c00082 | [6] |
| 2EK4 | -35.4 | 0.10 | 30 | 10.1021/acs.biochem.0c00082 | [6] |
| 2KE4 | -37.5 | 0.11 | 30 | 10.1021/acs.biochem.0c00082 | [6] |
| 3KE4 | -36.2 | 0.10 | 30 | 10.1021/acs.biochem.0c00082 | [6] |
| 4EK4 | -36.2 | 0.09 | 30 | 10.1021/acs.biochem.0c00082 | [6] |
| 4KE4 | -35.3 | 0.08 | 30 | 10.1021/acs.biochem.0c00082 | [6] |
| 5A1 | -34.3 | 0.08 | 39 | 10.1074/jbc.M401074200 | [7] |
| 6A1 | -34.1 | 0.09 | 46 | 10.1074/jbc.M401074200 | [7] |
| 7A1 | -34.7 | 0.08 | 53 | 10.1074/jbc.M401074200 | [7] |
| Lpp | -36.2 | 0.08 | 56 | 10.1021/bi048365%2B | [8] |
| PQLItet | -36.2 | 0.07 | 30 | 10.1039%2Fd2sc04479j | [9] |
| PQLL | -33.6 | 0.07 | 30 | 10.1039%2Fd2sc04479j | [9] |
| pqli3 | -28.7 | 0.08 | 23 | 10.1039%2Fd2sc04479j | [9] |
| pqll3 | -27.1 | 0.09 | 23 | 10.1039%2Fd2sc04479j | [9] |
| apCC_tetB | -33.8 | 0.09 | 30 | 10.1039%2Fd2sc04479j | [9] |
| apCC_tetA | -32.9 | 0.09 | 23 | 10.1039%2Fd2sc04479j | [9] |
| pqll_B | -36.8 | 0.09 | 30 | 10.1039%2Fd2sc04479j | [9] |
| huang2014 | -38.3 | 0.11 | 80 | 10.1126/science.1257481 | [10] |
| P1 | -14.0 | 0.06 | 4 | 10.1073/pnas.232591399 | [11] |
| P2 | -22.8 | 0.14 | 8 | 10.1073/pnas.232591399 | [11] |
| P3 | -29.5 | 0.10 | 11 | 10.1073/pnas.232591399 | [11] |
| 3 | -17.5 | 0.19 | 6 | 10.1021/ja0456003 | [12] |
| 13 | -18.0 | 0.19 | 6 | 10.1021/ja0456003 | [12] |
| chapman2004 | -29.9 | 0.03 | 7 | 10.1021/ja0466659 | [13] |
| dong2006 pep3 | -30.4 | 0.14 | 21 | 10.1021/bm050833n | [14] |

**Table S4. Best-fit parameters for the global fit to CD data using a linear spectroscopic model.** Parameter values are reported per peptide unit at 0°C and correspond to the mean of MCMC-obtained posterior distributions of model parameters (**Figure 6B**), along with their corresponding 2σ deviations.

| | |
|---|---|
| $\Delta G$ [kcal mol$^{-1}$ res$^{-1}$] | -0.23 ± 0.01 |
| $\Delta H$ [kcal mol$^{-1}$ res$^{-1}$] | -1.4 ± 0.1 |
| $\Delta C_p$ [kcal mol$^{-1}$ K$^{-1}$ res$^{-1}$] | 0.008 ± 0.002 |
| $v$ | 0.073 ± 0.004 |
| $k$ | 4.8 ± 1.3 |
| $[\theta]_{H\infty}$ [deg cm$^2$ dmol$^{-1}$ res$^{-1}$] | -42400 ± 1500 |
| $\partial[\theta]_{H\infty}/\partial T$ [deg cm$^2$ dmol$^{-1}$ °C$^{-1}$ res$^{-1}$] | 140 ± 40 |
| $[\theta]_C$ [deg cm$^2$ dmol$^{-1}$ res$^{-1}$] | 1550 ± 350 |
| $\partial[\theta]_C/\partial T$ [deg cm$^2$ dmol$^{-1}$ °C$^{-1}$ res$^{-1}$] | -40 ± 5 |

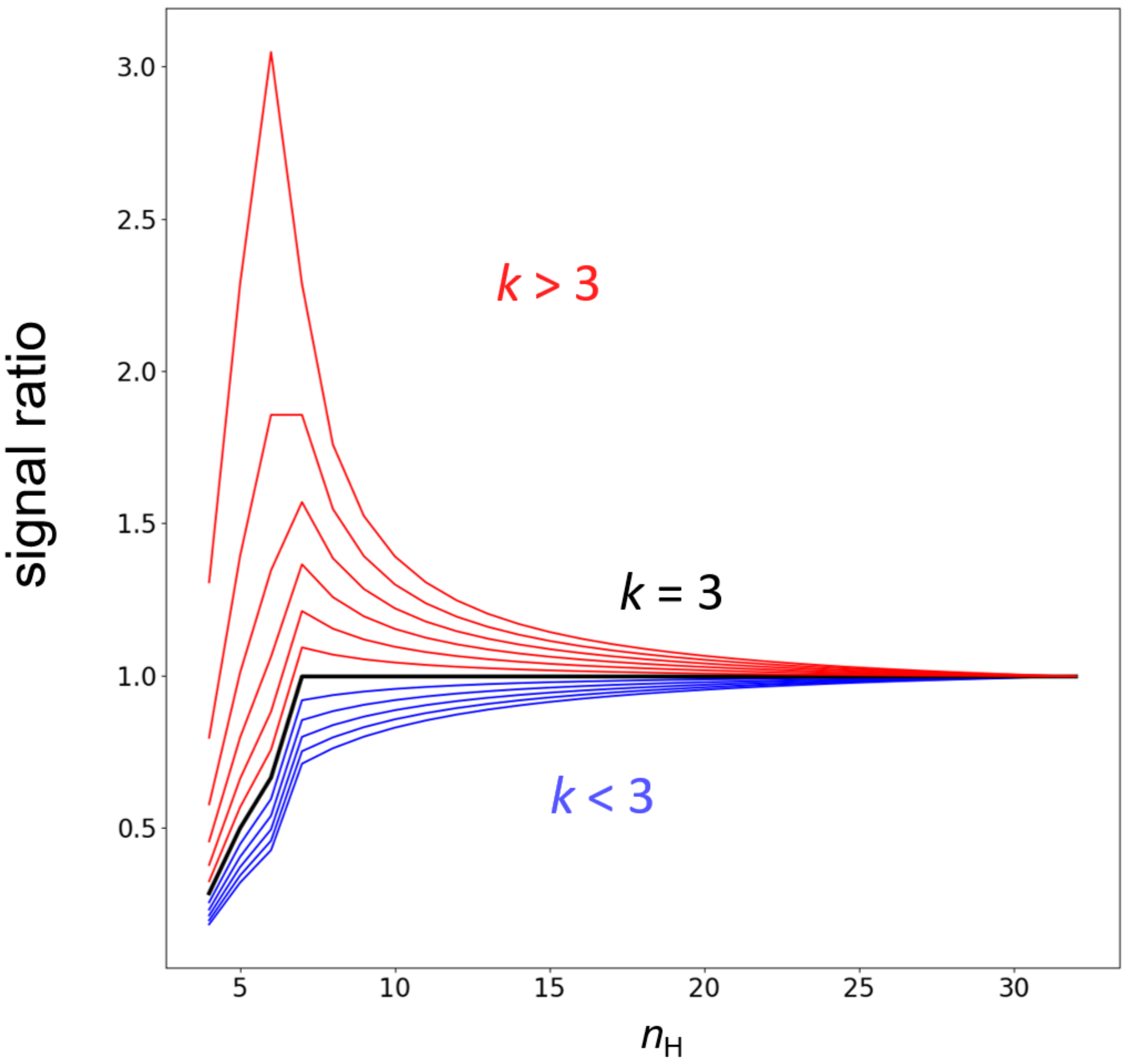


**Figure S1**. Ratio of predicted ellipticities $[\theta]_H$ for linear and dichroic model as a function of end-effect parameter *k*. For $k < 3$ (blue lines) CD signal is underestimated by linear model, while for $k > 3$ signals are overestimated compared to dichroic model, but eventually in both cases converge to common value at high $n_H$. In a special case when $k = 3$ both models give same result (except for short helices). The parameters used in calculation are same as in Figure 1.

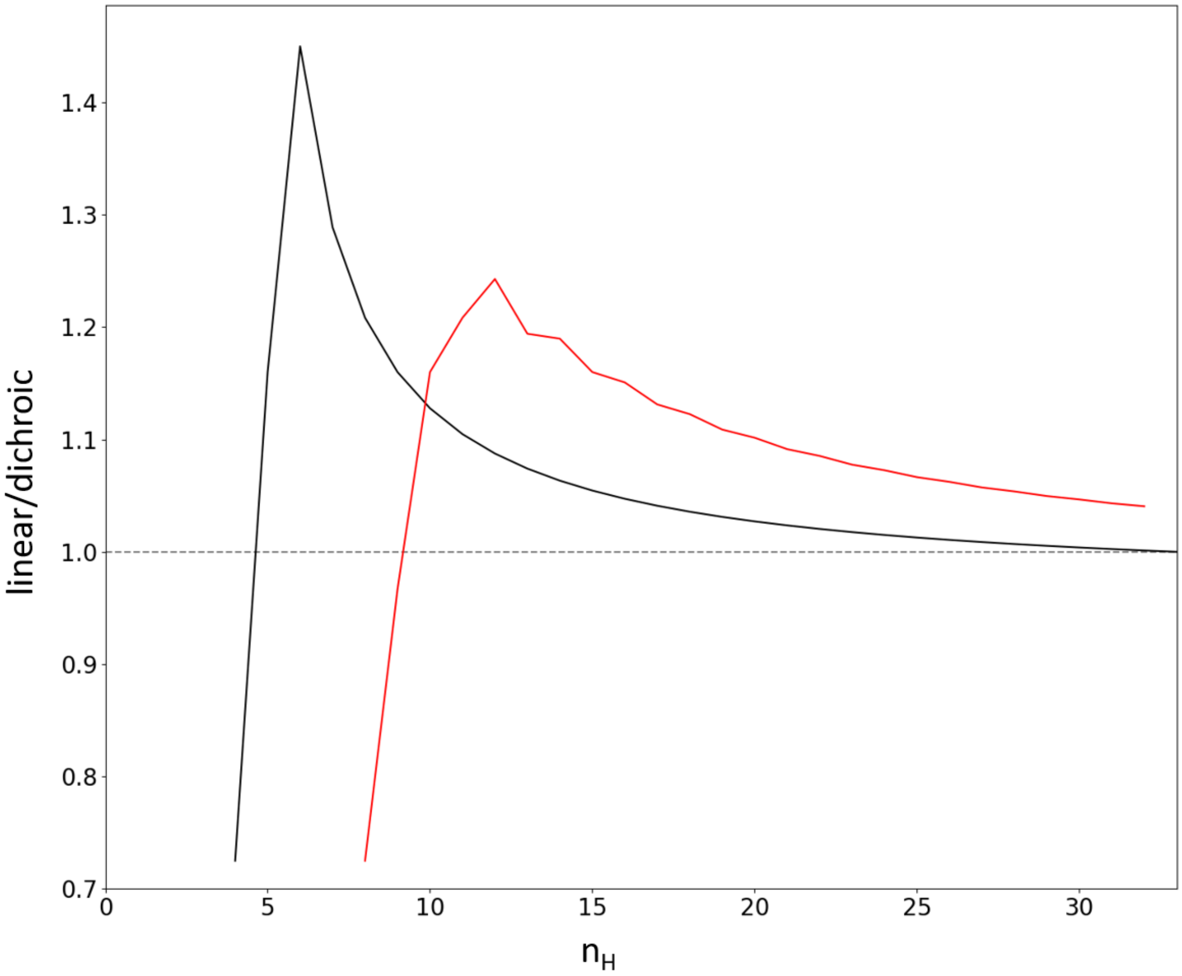


**Figure S2**. Ratio of ellipticities $[\theta]_H$ for linear vs. dichroic model for double-helix segments (red line). Ratio for single-helix is also shown for comparison (black line). For double helix segments signals do not converge even at high $n_H$ and linear model overpredicts CD signal across the full range of $n_H$ values. The parameters used in calculation are same as in Figure 1.

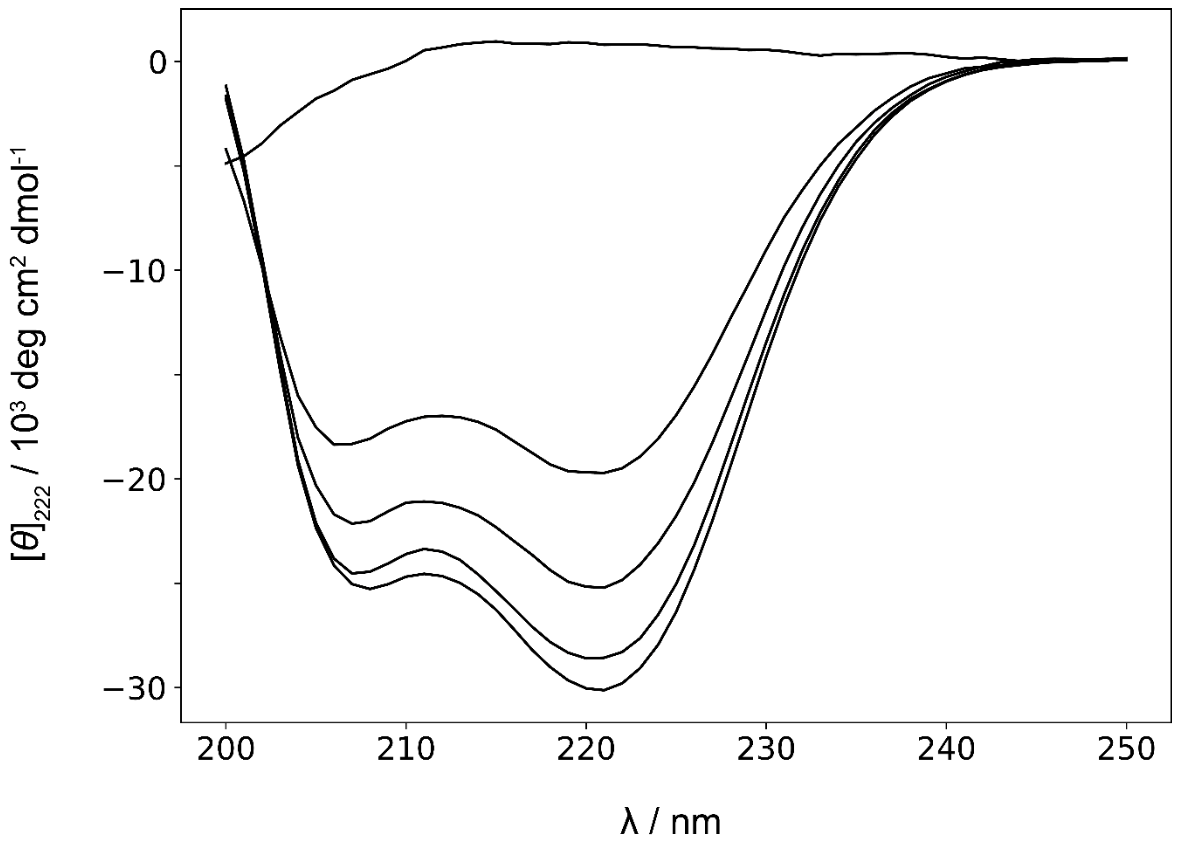


**Figure S3.** CD spectra of (AAKAA)$_n$-GY n=1,3,4,5,6 peptides at 0°C.

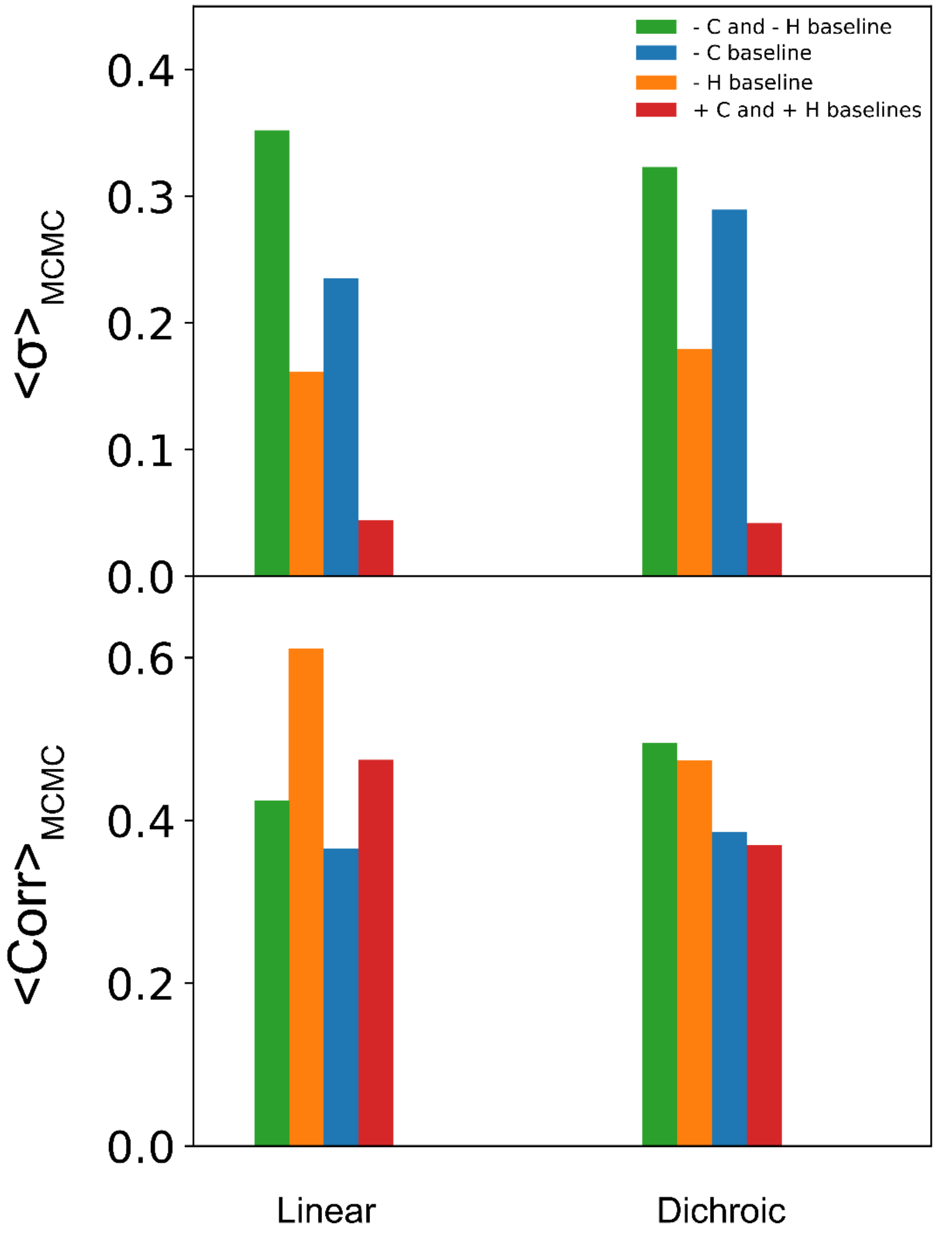


**Figure S4. Amount of information affects the uncertainty of determined model parameters.** Different fitting scenarios show that the additional data corresponding to the short peptide melt (AAKAA, termed "C baseline") and to all-helix dataset (termed "H baseline") constrain parameter values and results in more defined model. Upper panel shows average width (2σ) of parameters' posterior distributions obtained by performing MCMC (see **Methods**). Panel bellow shows average cross-correlations of model parameters obtained by analysis of MCMC corner plot (Spearman's rank correlation coefficient).

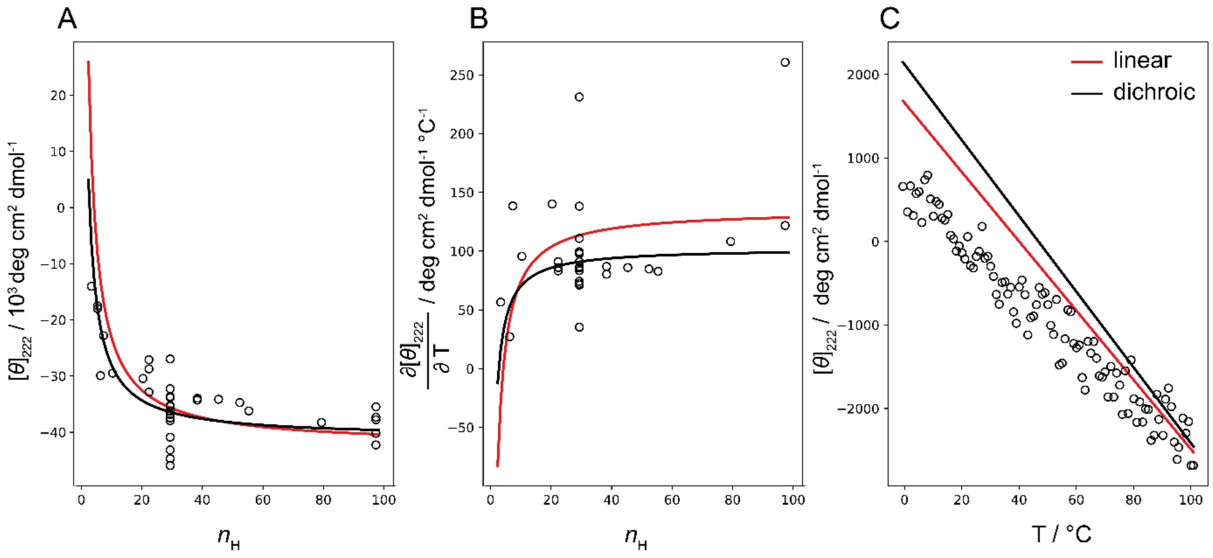


**Figure S5.** Spectroscopic baseline estimation with global fitting using linear or dichroic model. Red and black lines represent global-fit derived model functions describing spectroscopic baseline data with linear and dichroic model, respectively. Data is shown as symbols. A) Helix baseline parameter $[\theta]_H(n_{H,max})$ estimation using $[\theta]_{222}$ data for 39 all-helix systems at 0°C. B) Estimation of temperature-dependence of helix baseline ($\partial[\theta]_H(n_H)/\partial T$). C) Coil baseline parameters estimation ($[\theta]_C$ and ($\partial[\theta]_C/\partial T$).

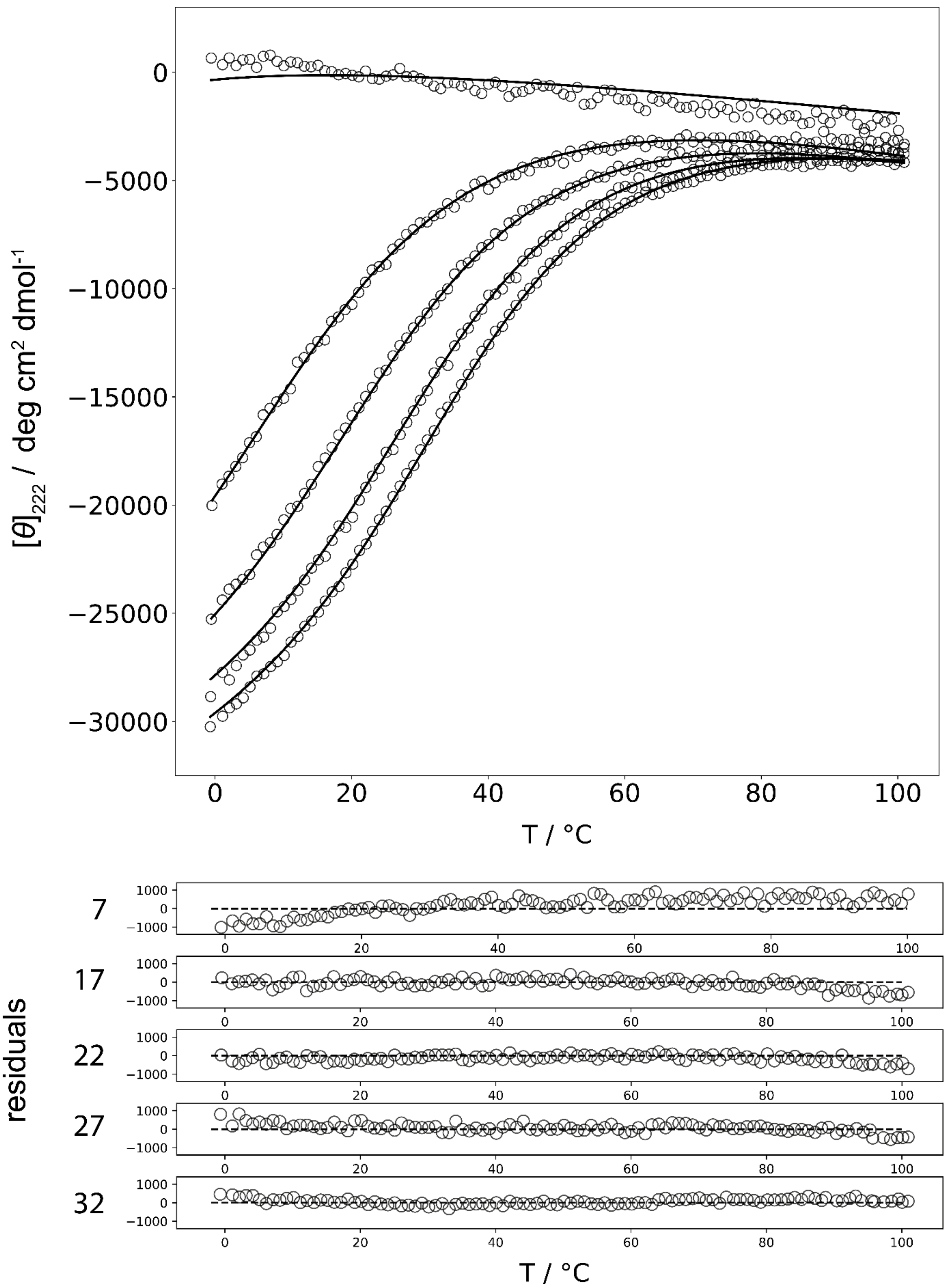


**Figure S6.** Global fit of CD data with the empirical model. In the upper panel data is shown as symbols and model function is shown with black full line. In the panel bellow residuals (model-data) are presented. As evident from the figure empirical model fails to describe high temperature data and also fails to establish appropriate coil baseline (data of shortest AAKAA peptide).

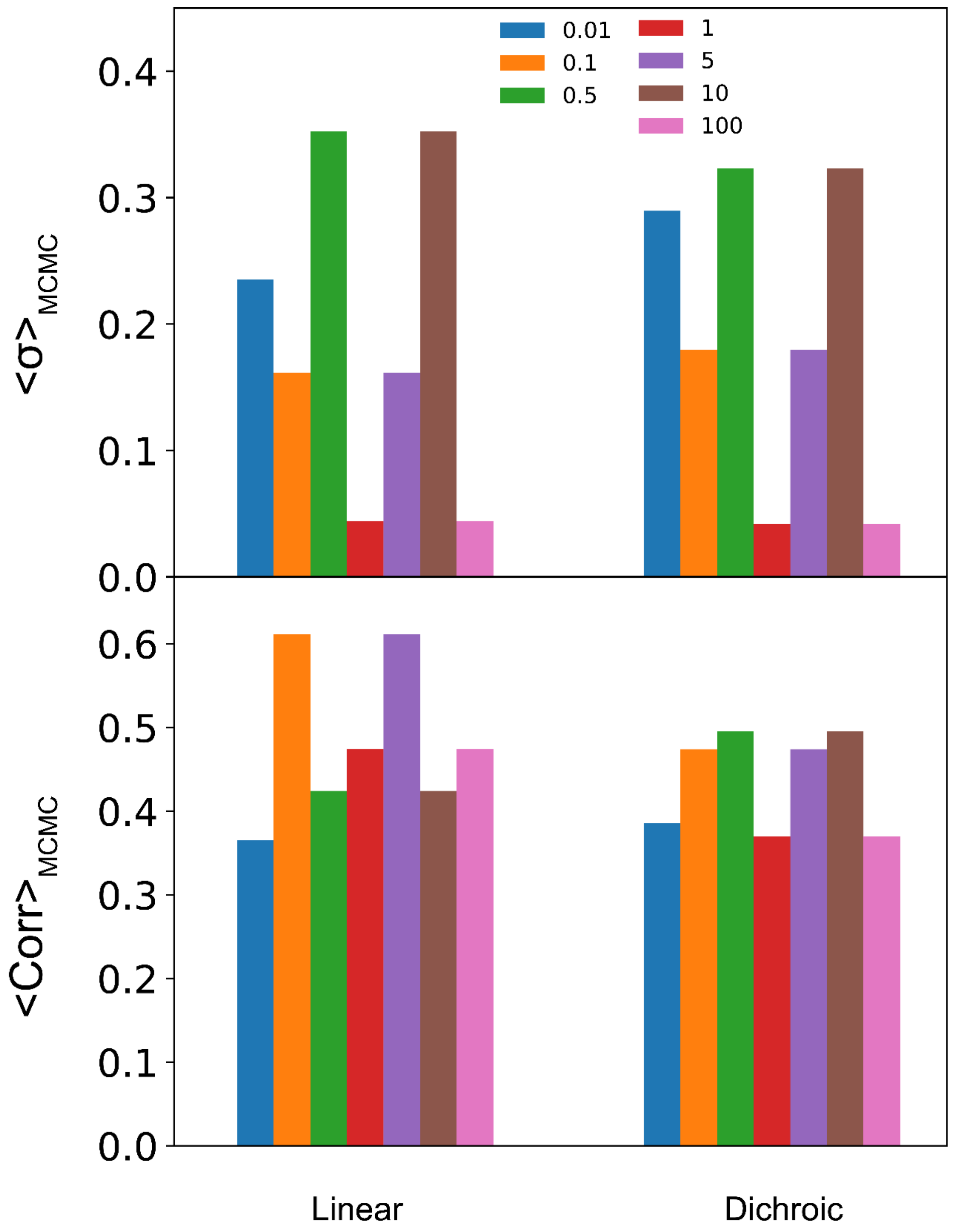


**Figure S7.** Weighted fitting of CD data with linear and dichroic model. Values of weights <1 give more attention to the CD melting $(AAKAA)_n$-GY data in the fitting process. On the other hand, weights >1 lead to fits that are more sound with all-helix baseline data (ignoring deviations in CD melting data). Upper panel shows average width (2σ) of parameters' posterior distributions as inferred by MCMC using different weights. Panel bellow shows average parameter cross-correlations obtained from MCMC corner plot (Spearman's rank correlation coefficient).

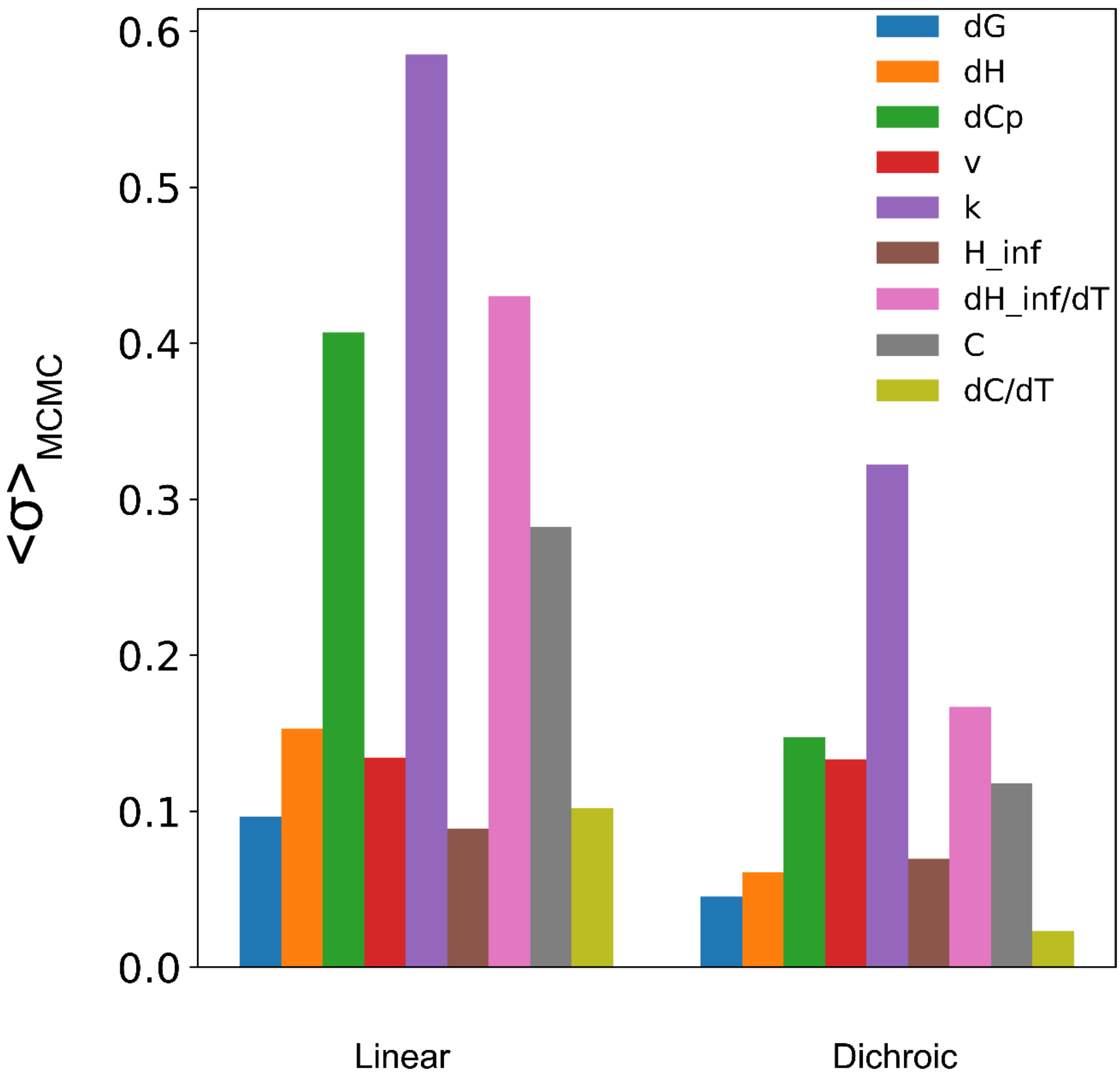


**Figure S8.** Best-fit parameter stability upon imposed weights on the data during fitting procedure (see **Figure S6**). Standard deviation of parameters' means of posterior distributions obtained via MCMC.

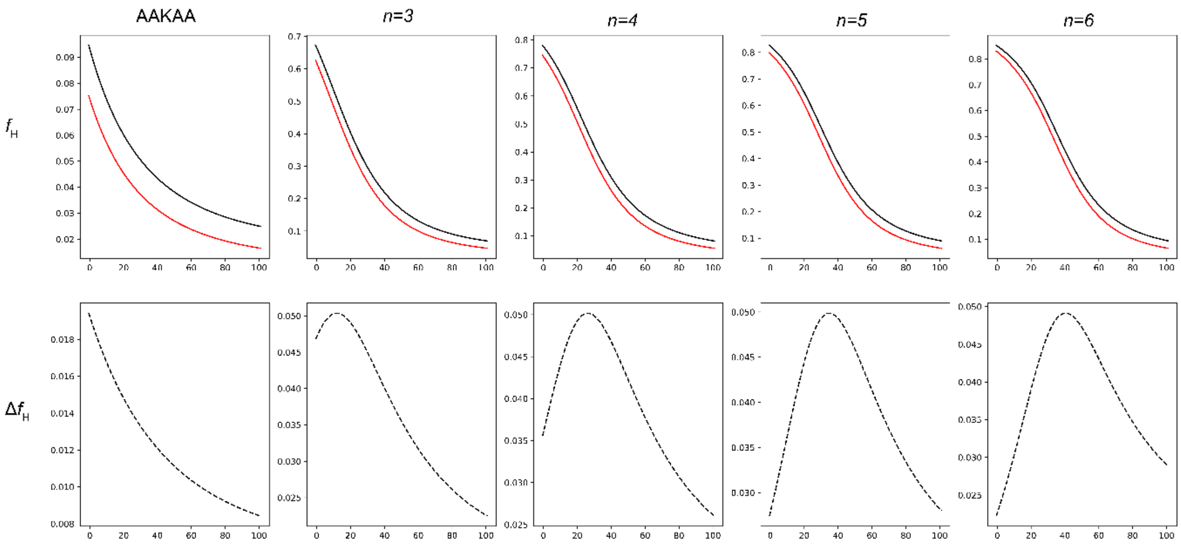


**Figure S9.** Helicity prediction as a function of temperature by linear (red) and dichroic model (black). Difference in helicities for different $(AAKAA)_n$-GY peptides are shown on the panels bellow.

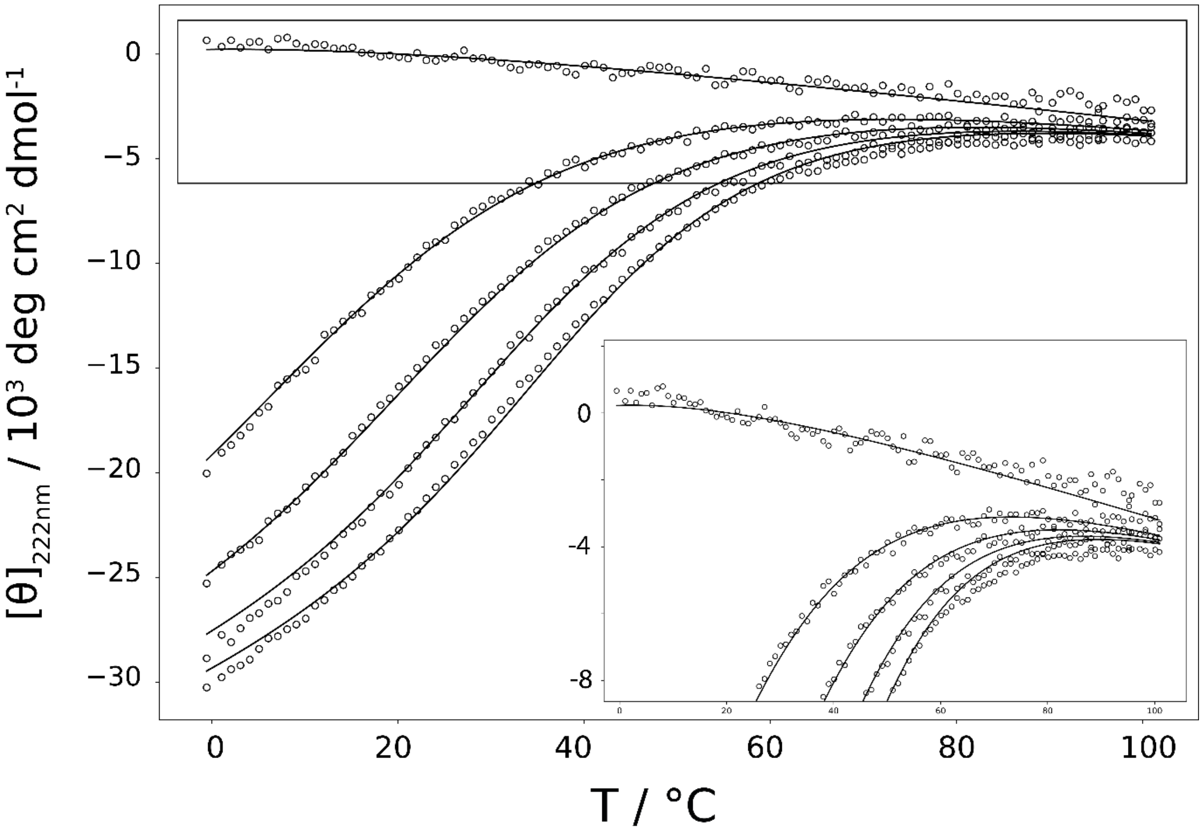


**Figure S10.** Global fit of dichroic model to CD data assuming $\Delta C_p = 0$ as a fixed parameter results in poorer description of the CD melting data. Deviation from the model prediction is especially high at higher temperatures and for shortest AAKAA peptide (inset).

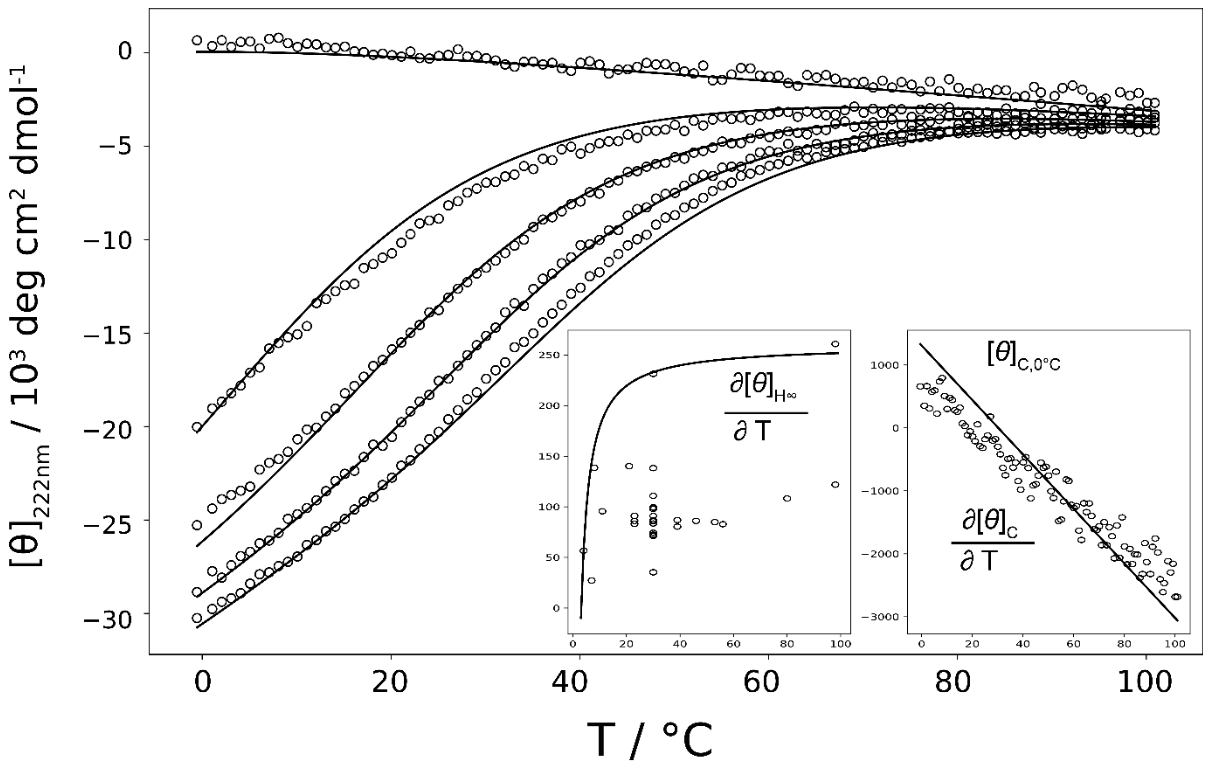


**Figure S11.** Global fit to CD data assuming $v$ = 0.048 as a fixed parameter results in poorer description of the CD melting data. Global description also fails to take into account spectroscopic baseline data (temperature dependence of helix ellipticity and coil baseline) – insets in the right bottom corner.

### References

[1] D. (Douglas) Poland and H. A. Scheraga, *Theory of helix-coil transitions in biopolymers; statistical mechanical theory of order-disorder transitions in biological macromolecules,*. Academic Press, 1970.

[2] I. Drobnak, H. Gradišar, A. Ljubetič, E. Merljak, and R. Jerala, “Modulation of Coiled-Coil Dimer Stability through Surface Residues while Preserving Pairing Specificity,” *J. Am. Chem. Soc.*, vol. 139, no. 24, pp. 8229–8236, Jun. 2017.

[3] M. Wolny *et al.*, “Characterization of long and stable de novo single alpha-helix domains provides novel insight into their stability,” *Sci. Rep.*, vol. 7, no. 1, pp. 1–14, Mar. 2017.

[4] G. G. Rhys *et al.*, “Navigating the Structural Landscape of de Novo α-Helical Bundles,” *J. Am. Chem. Soc.*, vol. 141, no. 22, pp. 8787–8797, Jun. 2019.

[5] N. R. Zaccai *et al.*, “A de novo peptide hexamer with a mutable channel,” *Nat. Chem. Biol.*, vol. 7, no. 12, pp. 935–941, Oct. 2011.

[6] C. L. Edgell, N. J. Savery, and D. N. Woolfson, “Robust de Novo-Designed Homotetrameric Coiled Coils,” *ACS Appl. Mater. Interfaces*, vol. 59, pp. 1087–1092, 2020.

[7] S. C. Kwok and R. S. Hodges, “Stabilizing and destabilizing clusters in the hydrophobic core of long two-stranded α-helical coiled-coils,” *J. Biol. Chem.*, vol. 279, no. 20, pp. 21576–21588, May 2004.

[8] ‡ Anatoly I. Dragan, § Sergey A. Potekhin, ∥ Andrei Sivolob, ⊥ and Min Lu, and ‡ Peter L. Privalov*, “Kinetics and Thermodynamics of the Unfolding and Refolding of the Three-Stranded α-Helical Coiled Coil, Lpp-56†,” 2004.

[9] E. A. Naudin *et al.*, “From peptides to proteins: coiled-coil tetramers to single-chain 4-helix bundles,” *Chem. Sci.*, vol. 13, no. 38, pp. 11330–11340, Sep. 2022.

[10] P. S. Huang *et al.*, “High thermodynamic stability of parametrically designed helical bundles,” *Science (80-. ).*, vol. 346, no. 6208, pp. 481–485, Oct. 2014.

[11] D. H. Chin, R. W. Woody, C. A. Rohl, and R. L. Baldwin, “Circular dichroism spectra of short, fixed-nucleus alanine helices,” *Proc. Natl. Acad. Sci. U. S. A.*, vol. 99, no. 24, pp. 15416–15421, Nov. 2002.

[12] N. E. Shepherd, H. N. Hoang, G. Abbenante, and D. P. Fairlie, “Single turn peptide alpha helices with exceptional stability in water,” *J. Am. Chem. Soc.*, vol. 127, no. 9, pp. 2974–2983, Mar. 2005.

[13] R. N. Chapman, G. Dimartino, and P. S. Arora, “A highly stable short α-helix constrained by a main-chain hydrogen-bond surrogate,” *J. Am. Chem. Soc.*, vol. 126, no. 39, pp. 12252–12253, Oct. 2004.

[14] H. Dong and J. D. Hartgerink, “Short homodimeric and heterodimeric coiled coils,” *Biomacromolecules*, vol. 7, no. 3, pp. 691–695, Mar. 2006.
